# MEF2A suppresses replicative stress responses that trigger DDX41-dependent IFN production

**DOI:** 10.1101/2022.09.15.508100

**Authors:** Julian R. Smith, Jack W. Dowling, Andrew Karp, Johannes Schwerk, Ram Savan, Adriana Forero

## Abstract

Interferons (IFN) are induced by sensing of self- and non-self DNA or genomic lesions by pathogen recognition receptors (PRR) that activate STING. These pathways must be kept in check by negative regulators to prevent unscheduled activation of IFN, which contributes to autoinflammation. Here we show that MEF2A as a novel negative regulator of inflammation that suppresses homeostatic induction of IFNs. Indeed, MEF2A deficiency results in the spontaneous production of type I IFN and robust downstream IFN-stimulated gene expression that coincided with a robust cellular antiviral state. Mechanistically, MEF2A depletion promoted the accumulation of R-loops that activate the DDX41/STING pathway. This pro-inflammatory pathway was dependent on ATR kinase activity, hallmark of the replicative stress response, was necessary for the activation of STING upon loss of MEF2A expression. Thus, our study connects MEF2A with protection from maladaptive type I IFN responses triggered by R-loop accumulation and links the DDX41-dependent activation of STING to the DNA damage response.

## INTRODUCTION

The detection of nucleic acids or DNA breaks by pattern recognition receptors (PRR) that promotes the formation of signalosomes that recruit Tank-binding kinase 1 (TBK1) to phosphorylate transcription factors, including as interferon regulatory factor 3 (IRF3). Phosphorylation of C-terminal serine residues of IRF3 allow for its dimerization and translocation to the nucleus where it transactivates type I interferon (IFNα/β) gene expression. Secreted IFNs can signal in an autocrine and paracrine manner through the type I IFN receptor to induce the phosphorylation of signal transducer and activator of transcription (STAT) factors, STAT1 and STAT2. The STAT dimers bind IRF9, forming the transcriptionally active complex, ISGF3, which binds to IFN-stimulated response elements (ISRE) positioned in hundreds of promoters that drive IFN-stimulated gene (ISG) expression (Chemudupati et al., 2019). The ISGs are critical effectors of the antiviral immune response and regulate various cellular processes to inhibit pathogen replication and dissemination. In addition, a subset of these ISGs is pro-inflammatory, promoting immune cell recruitment and coordinating tissue repair. Left unrestrained, IFN production can cause noxious tissue-damaging inflammation and pathogen-associated immunopathology. While the regulation of IFN responses is best characterized in the context of viral infection, accumulation and mis-localization of self-nucleic acids or the loss of nuclear genome integrity can also drive sterile IFN-mediated inflammation. These latter processes are the hallmark of autoinflammatory pathologies known as interferonopathies (Crow and Manel, 2015). Thus, it is critical to define the factors that protect organelle and genomic integrity, thereby preventing the excessive induction of type I IFN.

The major nucleic acid sensing pathways induced by the accumulation of mis-localized or damaged DNA converge on the adaptor molecule, Stimulator of Interferon Genes (STING). Three nucleic acid sensors have been characterized as upstream regulators of STING: cyclic GMP-AMP synthase (cGAS), IFN gamma inducible factor 16 (IFI16), and DEAD-box helicase 41 (DDX41). Cellular exposure to ionizing radiation can promote double-stranded DNA (dsDNA) breaks recognized by cGAS (Harding et al., 2017; Mackenzie et al., 2017), leading to the synthesis of the second messenger, 2’,3’-cyclic GMP-AMP (cGAMP), which promotes STING dimerization and activation. Genomic DNA breaks can also engage a non-canonical pathway that requires IFI16 and leads to cGAS-independent STING activation (Dunphy et al., 2018). Accumulation of immunostimulatory nucleic acids can also promote STING activation independent of damaged DNA. For example, deficiencies in the DNA exonuclease, TREX1, abrogates the clearance of cytoplasmic ssDNA, enhancing autoimmune responses in a STING-dependent manner (Stetson et al., 2008; Yang et al., 2007). Mutations in the RNase H2 exonuclease complex subunits, commonly associated with the autoinflammatory disorder Aicardi-Goutières syndrome (AGS), and induces STING-mediated inflammation (Crow et al., 2020) due to an accumulation of R-loops (Cristini et al., 2022). R-loops are triple-stranded RNA:DNA hybrid structures occurring naturally in ~5-10% of the genome (Brickner et al., 2022). In this regards, a subset of DEAD-box RNA helicases recognize unusual DNA:RNA structures, functioning to maintain genomic stability, but also serving as critical mediators of immunity (Andrisani et al., 2022; Cargill et al., 2021). One such helicase, DDX41, activates STING after recognition of dsDNA (Lee et al., 2015; Singh et al., 2022; Zhang et al., 2011) and RNA:DNA transcripts generated by retroviral infection (Stavrou et al., 2018). Recent studies have shown that DDX41 is also recruited to R-loops, promoting their resolution and preventing DNA. However, a role, if any, for DDX41-dependent processes in the induction of IFN-mediated inflammation after R-loop accumulation remains to be determined.

A critical first step in repair of DNA breaks is activation of the DNA damage response (DDR) kinases Ataxia–telangiectasia mutated (ATM), ATM and RAD3-related (ATR), and DNA-dependent protein kinase catalytic subunit (DNA-PKcs). In addition to coordinating cell cycle arrest and repair, DDR kinases regulate host inflammatory responses associated to genotoxic stress (Blackford and Jackson, 2017; Forero et al., 2014), cytosolic DNA accumulation (Burleigh et al., 2020; Ferguson et al., 2012), and DNA virus infection (Justice and Cristea, 2022). Amongst these kinases, ATR is the central regulator of the replicative stress response (RSR) that occurs under conditions associated with stalled replication fork progression, such as nucleotide pool depletion and R-loop accumulation. ATR phosphorylates substrates that mediate cell cycle arrest to prevent chromosomal breaks (Ragu et al., 2020; Saldivar et al., 2017), promotes R-loop resolution, and proper chromosome segregation (Hodroj et al., 2017; Kabeche et al., 2018; Matos et al., 2020). While ATM and DNA-PKcs have been linked to IRF3 activation and IFN-driven inflammatory responses, the mechanisms by which ATR contributes to IFN production are less clear.

Recently, the myocyte enhancing factor 2 (MEF2) transcription factors have emerged as potential regulators of inflammation (Cilenti et al., 2021; Deczkowska et al., 2017; Lu et al., 2021; Xue et al., 2021). The MEF2 family is composed of 4 evolutionarily conserved proteins, MEF2A-D, that temporally coordinate cardiac and neuronal development (Chen et al., 2017). MEF2 proteins have important homeostatic roles as mice deficient in these factors display embryonic lethality or early post-natal lethality (Lin et al., 1997; Naya et al., 2002). Our study focuses on MEF2A, as conflicting studies have examined the association of loss of function (LOF) mutations with an enhanced risk for adverse coronary artery disease outcomes. IFN induction following cardiac injury has been proposed to shape tissue remodeling and contribute to worsened outcomes (King et al., 2017). Moreover, MEF2A is the least characterized immune regulator in this family. Using molecular approaches, we show that acute depletion of MEF2A can drive spontaneous IFN production and a downstream cellular antiviral state. We demonstrate that MEF2A expression can protect genomic integrity by preventing the excessive accumulation of R-loops. This increase in RNA:DNA hybrid formation drives DDX41/STING-mediated IFN responses. Lastly, we show that ATR kinase activity is required to support STING activation in this context. These findings position MEF2A as a positive regulator of genomic stability which protects from unscheduled IFN-driven inflammation across various cell types.

## RESULTS

### Loss of MEF2A leads to spontaneous IFN production

Given the potential connection between MEF2 proteins and inflammation, we examined the impact of one MEF2 family member on the transcriptional program in myocytes, a cellular target of inflammation caused by viral infections. MEF2A is the most abundantly expressed among the MEF2 family of transcription factors in neonatal myocytes, but its role in adult cardiomyocytes is poorly understood. As such, we used small interfering (siRNA) to transiently deplete *MEF2A* in the human cardiomyocyte cell line, AC16. Transient depletion of *MEF2A* allowed us capture acute gene expression changes while circumventing possible functional compensation by other MEF2 paralogs, as previously described in knock-out cells (Majidi et al., 2019). Genome-wide transcriptional signatures captured by RNA-sequencing revealed robust differentially expressed (DE) transcripts, in which 519 genes were significantly upregulated and 201 genes were significantly downregulated in *MEF2A*-targeted cells compared to cells transfected with non-targeting control siRNA (**Figure 1A, Supplementary Table 1**). Gene ontology analysis identified enriched biological functions amongst the upregulated genes following MEF2A depletion. We observed significant enrichment in pathways associated with innate immune inflammation and type I IFN responses amongst the upregulated genes (**Figure 1B**). This was consistent with an increase in the expression of *IFNB1* and *CXCL10* mRNA in MEF2A depleted cells (**Figure 1C**). To validate the reproducibility of *MEF2A*-targeting in driving IFN induction we tested two additional MEF2A-targeting siRNAs. We found that in all cases, *MEF2A* silencing led to the phosphorylation of the IFN responsive transcription factor STAT1 at tyrosine 701 (Y701) relative to non-targeting siRNA transfected cells **(Figure S1A**).

**Figure 1.**
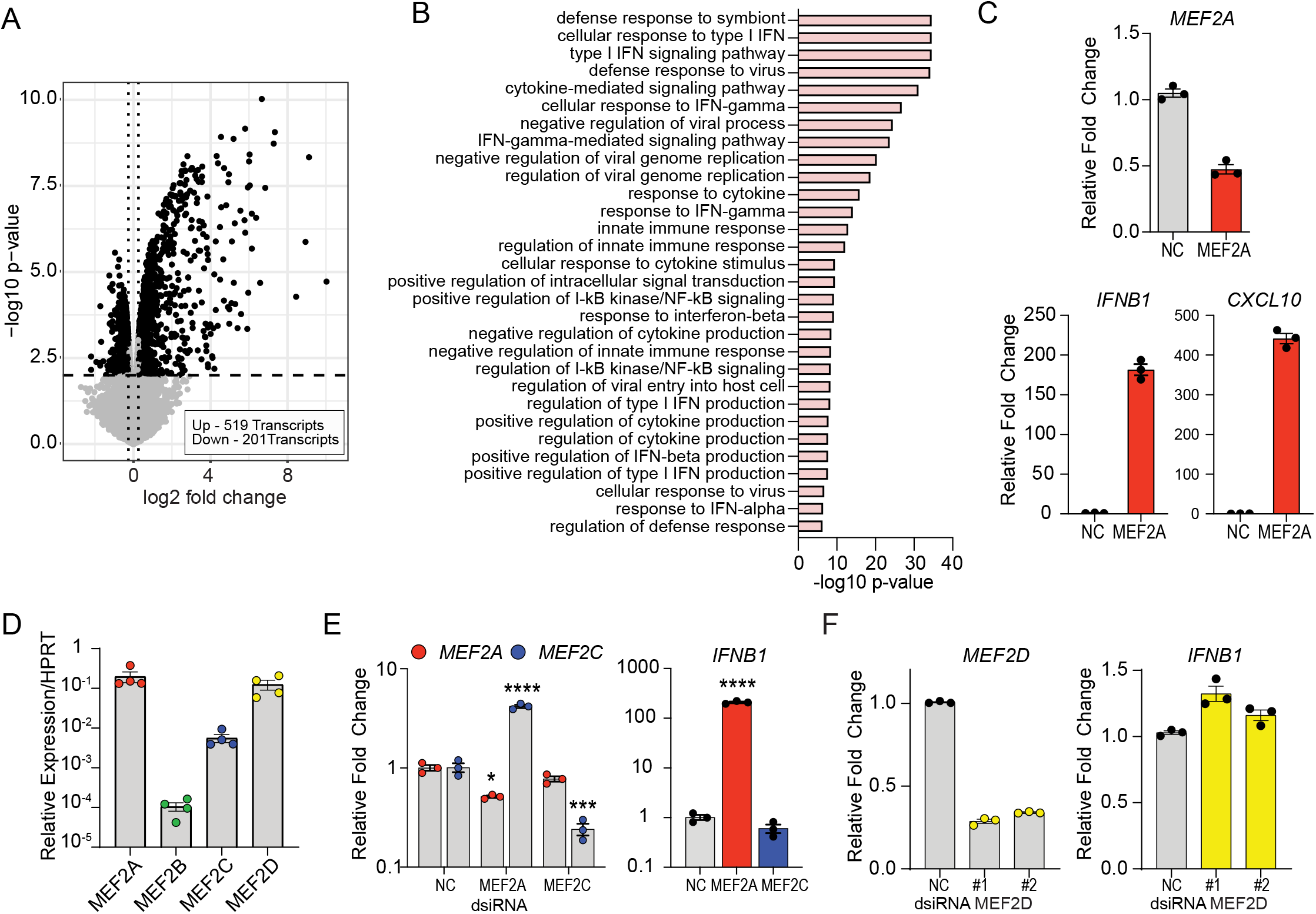
MEF2A silencing is sufficient to induce innate immune inflammatory responses. (A) Loss of MEF2A drives a robust change in transcriptional expression. Volcano plot of differentially expressed genes (LFC |0.26|; adj p-value 0.01) following knockdown (KD) of *MEF2A* with dsiRNA. 519 transcripts were significantly upregulated, and 201 transcripts were downregulated (black) relative to non-targeting control (NC) transfected cells. Each dot represents a unique transcript, dashed lines indicate threshold of significance. Non-significant changes in gene expression are highlighted in grey. (B) Gene Ontology enrichment analysis demonstrates that loss of MEF2A promotes changes in genes involved in innate immune antiviral pathways. Bar graph represents the top 30 GO biological processes enriched amongst the genes upregulated by *MEF2A* depletion. (C) Loss of MEF2A promotes the expression of *IFNB1* and *CXCL10*. Relative expression of *MEF2A, IFNB1*, and *CXCL10* mRNA following MEF2A depletion. Relative expression was normalized to NC transfected cells (value of 1) and normalized to *HPRT1*. Data is representative of the average of three individual replicates and error bars represent SEM. (D) Relative quantity of endogenous MEF2 gene expression in AC16 cells. Expression of MEF2 transcripts was calculated relative to *HPRT1*. Bars represent the average expression across 4 independent experiments and error bars represent SEM. (E-F) The induction of IFN is specific to *MEF2A* silencing. (E) Bar graphs represent relative expression of *MEF2A* (red) and *MEF2C* (blue) mRNA following dsiRNA transfection (left). Expression of *IFNB1* mRNA following siRNA-mediated depletion of *MEF2A* (red) or *MEF2C* (blue). Relative fold change was calculated relative to NC transfected cells and normalized to *HPRT1* (value 1). (F) Bar graphs represent the average relative expression of *MEF2D* (yellow, left) and *IFNB1* (right) mRNA following transfection of AC16 cells with two distinct MEF2D targeting dsiRNA or NC. Relative fold change in expression was calculated relative to *HPRT1* and NC transfected cells (value 1). Experiments represent average of 3 individual experiments. Asterisks indicate **** p ≤ 0.0001, *** p ≤ 0.001 and * p < 0.05 as determined by one-way ANOVA.

Consistent with the expression of MEF2 transcription factors in left ventricular adult human cardiomyocytes (GTEX) **(Figure S1B**), AC16 cells express high levels of *MEF2A* and *MEF2D*, moderate levels of *MEF2C*, and low levels of *MEF2B* mRNA (**Figure 1D).** To determine whether a single MEF2 factor (Desjardins and Naya, 2016) was responsible for IFN induction, we independently targeted MEF2A, MEF2C and MEF2D in AC16 cells. Although depletion of *MEF2A* resulted in an increase in *MEF2C* transcript (**Figure 1E, left**), the depletion of *MEF2C* did not induce *IFNB1* mRNA expression relative to non-targeting control transfected cells (**Figure 1E, right**). Similarly, the silencing of *MEF2D* did not lead to significant changes in *IFNB1* mRNA expression relative to control transfected cells (**Figure 1F**). We recapitulated this phenotype in immortalized human fibroblasts, which have similar MEF2 gene expression profiles to myocytes (**Figure S1C**). Silencing of *MEF2A* was sufficient to induce both *IFNB1* mRNA and that of ISGs such as *CXCL10* (**Figure S1D, left**). Finally, we observed similar upregulation of *IFNB1* and *CXCL10* mRNA in human macrophage-like cells, THP-1, induced by the knockdown of *MEF2A* expression (**Figure S1D, right**). Together, these results suggest that MEF2A, but not other MEF2 protein family members, is a negative regulator of spurious type I IFN transcription cross multiple human cell types.

### The type I IFN responses predominate upon MEF2A depletion

Having observed an enrichment of genes involved in IFN responses, we addressed whether this gene signature was associated with resistance to viral infection. *MEF2A* silencing rendered cells refractory to the cytolytic effects of vesicular stomatitis virus (VSV) infection compared to control cells as determined by crystal violet uptake assay (**Figure 2A**). Similarly, loss of MEF2A conferred AC16 cells with protection against the cardiotropic enterovirus, Coxsackievirus B3 (CVB3), which exhibited a 75% reduction in viral RNA accumulation relative to non-targeted cells (**Figure 2B**). Previously, we have demonstrated that such decreases in viral RNA correspond to a significant attenuation in infectious virus production (Soveg et al., 2021). These data suggest that MEF2A loss leads to the secretion of bioactive IFN that can establish a cellular antiviral state.

**Figure 2.**
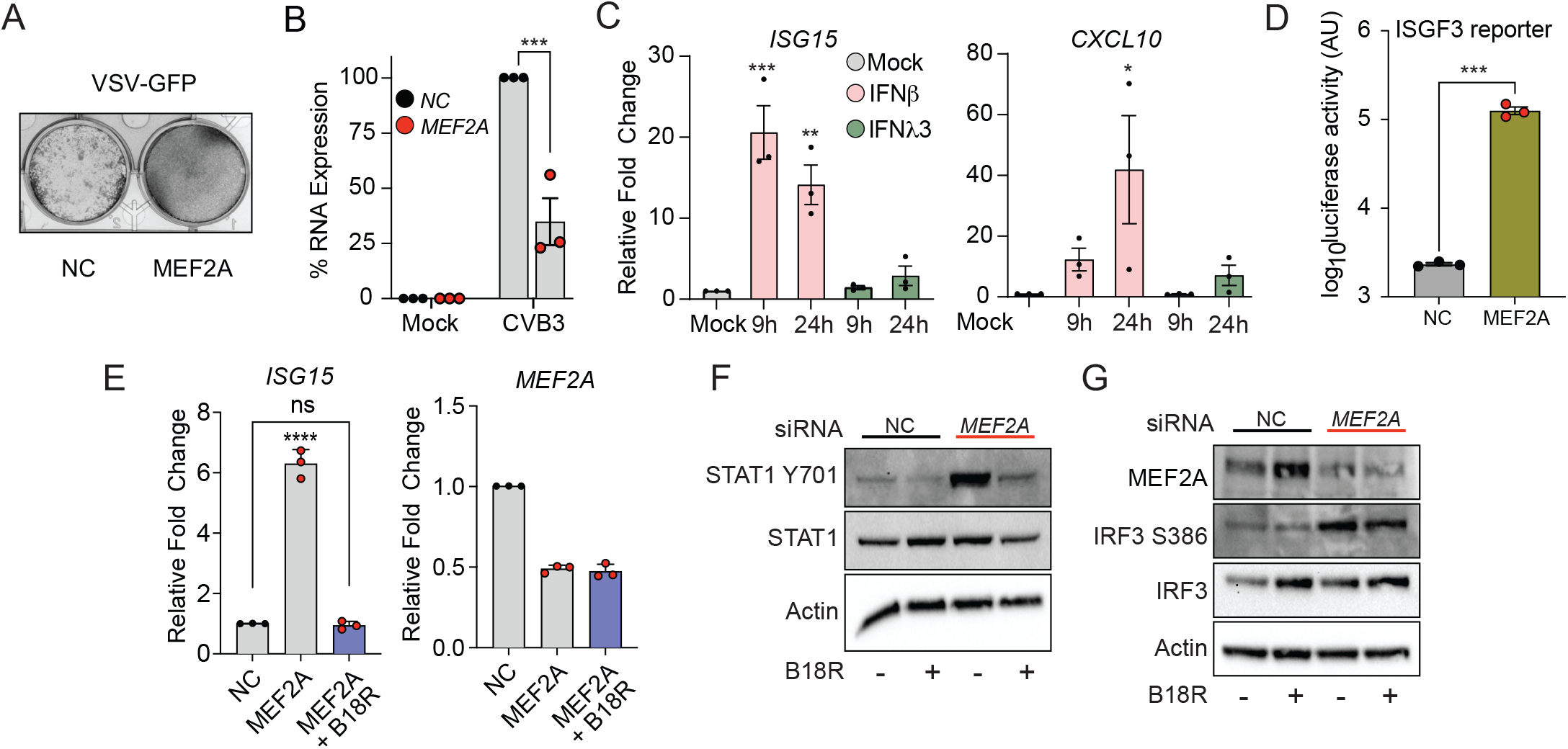
Silencing of MEF2A induced a type I IFN driven antiviral state. (A) Loss of *MEF2A* protects cells against the cytopathic effect of vesicular stomatitis virus (VSV). AC16 cell monolayers transfected with non-targeting control (NC) or *MEF2A*-targeting dsiRNA 24 hours (h) prior to infection with VSV-GFP. Crystal violet uptake was measured 24 h post VSV infection. (B) *MEF2A* loss protects against cardiotropic virus infections. Bar graphs represent percent reduction of coxsackie virus B (CVB3) viral RNA expression in *MEF2A*-depleted cells (red) relative to NC-transfected cells (black) across 3 independent experiments. Error bars represent SEM. (C) Responsiveness of cardiac muscle cells to exogenous recombinant IFN stimulation. AC16 cells were stimulated with 25 IU/ml IFNβ (pink) or 100 ng/ml IFNλ3 (green) for the indicated times prior to total RNA harvest. Relative expression of *ISG15* and *CXCL10* mRNA was measured relative to *HPRT1* and normalized to mock-treated cells (value 1). Bar graphs represent average of 3 independent experiments and error bars represent SEM. *** p ≤ 0.001, ** p ≤ 0.01, * p < 0.05 as determined by two-way ANOVA. (D) Secretion of IFN from AC16 cells following dsiRNA transfection. Bar graphs represent average *Gaussia* luciferase activity from 5xISGF3-Gluc Huh7 reporter cells stimulated with supernatants derived from AC16 cells transfected with either NC (grey bar) or *MEF2A* (gold bar) targeting dsiRNA. Data represents 3 independent experiments and error bars represent SEM. *** p ≤ 0.001, as determined by students t-test. (E) Blockade of secreted type I IFN inhibits ISGs following MEF2A-depletion. Type I IFN was inhibited by pre-treatment of AC16 cells with 1 μg/mL of recombinant B18R prior to transfection with NC or *MEF2A*-targeting dsiRNA. Expression of *ISG15* (left) or *MEF2A* (right) mRNA was quantified relative to *HPRT1* and normalized to NC-transfected cells (value of 1). Bar graphs represent average of 3 independent experiments, error bars represent SEM and **** represent p ≤ 0.0001 as determined by one-way ANOVA. (F) AC16 cells were pre-treated with B18R as previously described and transfected with dsiRNA. Activation of IFN signaling was probed by measuring STAT1 (Y701) phosphorylation and total STAT1 by western blot. Total Actin protein expression served as loading control. (G) IRF3 activation precedes IFN responses. AC16 cells were treated as mentioned above and the levels of phosphorylated IRF3 (S386), total IRF3, and Actin were measured by western blot analysis.

Cell intrinsic antiviral responses are coordinated by both type I and III IFNs. Both IFN families can promote the activation of ISGF3 to promote transcriptional induction of ISG expression (Kessler et al., 1988). As we observed both type I and III IFN gene mRNA induction after MEF2A depletion **(Supplementary Table 1)** we asked whether the secretion of IFN drove inflammation following *MEF2A* silencing. We examined whether human cardiomyocytes could respond to exogenous recombinant type I (IFNβ) and III (IFNλ3) IFN stimulation by measuring the induction of ISG mRNA after IFN treatment. Treatment with recombinant IFNβ induced significant expression of *ISG15* and *CXCL10* mRNAs at 9h and 24 h post stimulation (**Figure 2C**). In contrast, IFNλ3 stimulation did not induce the expression of either of these two ISGs (**Figure 2C**). We engineered a cell-based luciferase reporter assay to monitor ISGF3-mediated transcriptional activation in response to secreted IFN in Huh7 cells, which respond to exogenous IFN stimulation (**Figure S2A**). Transfer of supernatants from MEF2A depleted muscle cells led to a significant increase in luciferase activity relative to treatment with supernatants derived from non-targeted control transfected cells (**Figure 2D**). We then confirmed the predominant inflammatory role for type I IFN by treating AC16 cells with the pan-type I IFN inhibitor, B18R (Symons et al., 1995). Pretreatment with B18R decreased *ISG15* mRNA induction post *MEF2A* silencing (**Figure 2E**). This was concomitant with decreased STAT1 phosphorylation (Y701) in B18R pre-treated cells relative to vehicle treatment in response to *MEF2A* depletion (**Figure 2F**). Of note, levels of phosphorylated STAT1 induced by silencing of MEF2A were comparable to those induced by IFNβ stimulation, both of which enhanced IRF3 activation, as determined by serine 386 phosphorylation, and increased IRF1 protein expression (**Figure S2B**). Importantly, IRF3 activation was unaffected by type I IFN blockade with B18R (**Figure 2G**), suggesting that decreases in MEF2A promote signaling transduction cascades upstream IRF3 to drive IFN expression (**Figure 1A**). Together, these data suggest that IFN is secreted after MEF2A silencing and the establishment of the antiviral state in cardiomyocytes depended sole on type I IFN responses.

### Inflammatory responses in MEF2A depleted cells are STING-dependent

To further define signaling mechanisms for IFN responses in the wake of MEF2A deficiency, we first confirmed that IRF3 was indispensable for this process. For this purpose, we generated *IRF3*-deficient cardiomyocytes using CRISPR-Cas9 genome editing technologies. Deletion of IRF3 abrogated the activation of STAT1 in response to *MEF2A* silencing (**Figure 3A**), suggesting minimal compensation from additional IRFs, such as IRF1 or IRF7, which that can transactivate IFN genes (Forero et al., 2014; Miyamoto et al., 1988; Suschak et al., 2016) and are induced by MEF2A depletion (**Figure S2B**). IRF3 activation is mediated by PRRs that can sense nucleic acids, and converge on adaptor molecules such as MAVS (RNA sensing) and STING (DNA sensing) (Ablasser and Hur, 2020). We confirmed that both RNA and DNA sensing pathways were functional in our cellular model by measuring the induction of *IFNB1* (**Figure 3B**), *IFNL1* and *CXCL10* mRNAs (**Figure S3A**) following the transfection of cells with either a specific RIG-I agonist (Saito et al., 2008) or calf-thymus DNA (Ishikawa et al., 2009). In addition, the cardiomyocytes responded to TLR3 activation with poly(I:C) or infection with Sendai virus (SeV) (**Figure S3B**), which activates MAVS-dependent responses through RIG-I and MDA5 (Yount et al., 2008).

**Figure 3.**
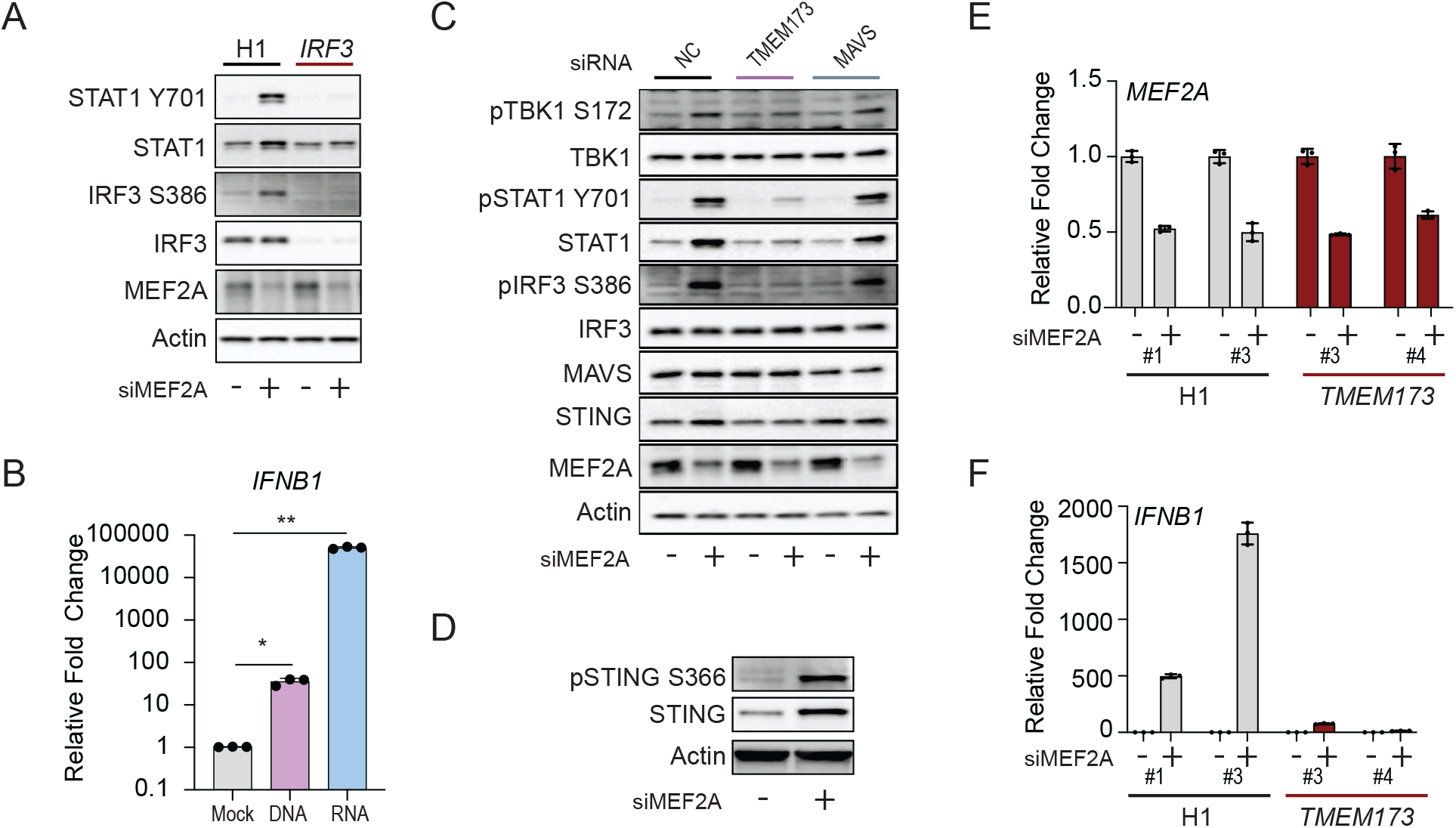
STING expression is necessary for the induction of IFN-mediated inflammation post MEF2A silencing. (A) Deletion of IRF3 abrogates induction of IFN-responses following *MEF2A* knockdown. AC16 cells stably transduced with non-targeting single-guide RNA (sgRNA) (H1) or IRF3-targeting sgRNA (IRF3) were transfected with NC or *MEF2A* targeting dsiRNA for 24 h. Analysis of protein expression analysis of IRF3 (S386) and STAT1 phosphorylation (Y701) by western blot. Total IRF3, STAT1, MEF2A, and Actin protein expression were also measured as controls. (B) AC16 have intact RNA and DNA sensing pathways. AC16 were transfected with calf-thymus DNA (DNA, purple) or HCV PAMP (RNA, blue) for 6 h prior to total RNA harvest. Bar graphs represent average relative fold changes in mRNA levels were calculated relative to *HPRT1* and normalized to mock-transfected cells (value 1). Bar graph represents 3 independent experiments and error bars represent SEM. ** p ≤ 0.01 and * p < 0.05 as determined by one-way ANOVA. (C) Signal transduction through STING, but not MAVS, is required for IFN induction after the loss of MEF2A. AC16 cells were transfected with dsiRNA targeting *MAVS, TMEM173* (STING), or NC. After 24 h, cells were transfected with siRNA targeting *MEF2A* or NC for an additional 24 h. Whole cell protein lysates were probed for TBK1 (S172), IRF3 (S386) and STAT1 (Y701) phosphorylation. Total protein expression of TBK1, STAT1, IRF3, MAVS, STING, MEF2A, and Actin were measured as controls. Western blot representative of 3 independent experiments. (D) Loss of MEF2A leads to the activation of STING. AC15 cells were transfected with NC or MEF2A targeting siRNA for 24 h. Whole cell lysate was subjected to protein expression analysis for STING (S366) phosphorylation, total STING and Actin. (E and F) STING deletion abrogates IFN induction following MEF2A knockdown. (E) AC16 H1 and *TMEM173* (STING) KO cells were targeted with dsiRNA against *MEF2A*. Bar graphs represent the average relative MEF2A mRNA expression relative to *HPRT1* expression and NC transfected cells (value 1) across 2 individual experiments. Numbers indicate clonal cell lines. (F) Relative expression of *IFNB1* mRNA induced by *MEF2A* silencing. Bar graphs represent relative fold changes in expression as normalized to *HPRT1* expression and NC transfected cells (value 1) across 2 individual experiments. Numbers indicate clonal cell lines.

To discern which of these nucleic acid sensing pathways is engaged in response to *MEF2A* silencing, we depleted either STING (*TMEM173*) or *MAVS* prior to targeting *MEF2A* expression. We found STING, but not MAVS, was necessary to promote the activation of TBK1 and IRF3 phosphorylation upstream of type I IFN induction. Similarly, we observed that STAT1 phosphorylation following *MEF2A* depletion (**Figure 3C**) was muted only in cells deficient for STING. Silencing of *MAVS* did not abrogate STAT1 phosphorylation relative to that observed in control cells (**Figure 3C**). Further, STING phosphorylation at serine 366 (Liu et al., 2015) was observed upon *MEF2A* depletion (**Figure 3D**). Next, we generated STING knockout cells by targeting *TMEM173* using CRISPR-Cas9 genome editing (**Figure S3C**). In these cells, silencing of *MEF2A* (**Figure 3E**), did not promote the accumulation of *IFNB1* mRNA (**Figure 3F**). Together, these data indicate that MEF2A mitigates the accumulation of DNA ligands that drive STING activation, and could promote spontaneous IFN-mediated inflammation.

### The loss of MEF2A compromises genomic integrity

The disruption of cellular homeostasis can induce STING-mediated inflammatory responses. Both the loss of nuclear DNA integrity and the accumulation of endoplasmic reticulum (ER) stress have been associated with STING-mediated inflammation in the absence of pathogen infection (Petrasek et al., 2013). To determine whether MEF2A is necessary to sustain either DNA integrity or ER function, we silenced MEF2A expression in AC16 cells, and measured DNA damage or ER stress response markers. Loss of MEF2A led to an accumulation of phosphorylated histone 2AX (γH2A.X), a marker of DNA damage (**Figure 4A**). Similar γH2A.X increases were observed following treatment with the topoisomerase II inhibitor, etoposide, known to elicit genotoxic stress (**Figure 4A**). However, neither the loss of *MEF2A* nor treatment with etoposide led to an accumulation of ATF4 or enhanced splicing of XBP1 (sXBP1), hallmarks of the ER stress response. As expected, treatment with thapsigargin, a sarco/ER calcium ATPase (SERCA) inhibitor, led to both increases in ATF4 protein expression and XBP1 splicing (**Supplementary Figure 4A**). We conclude that loss of MEF2A results in the accumulation of γH2A.X and DNA damage, which activates STING and type I IFN induction.

**Figure 4.**
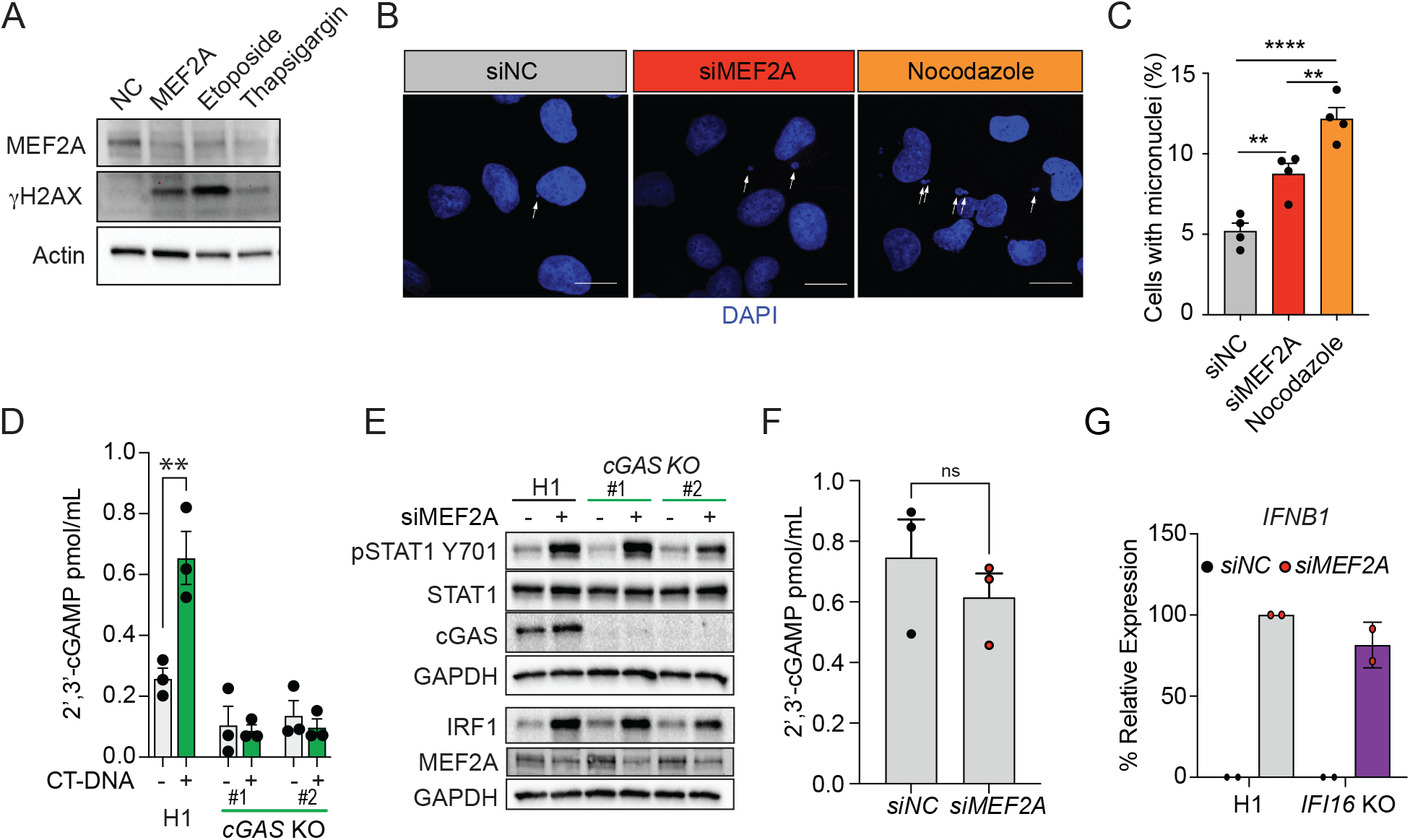
The loss of MEF2A compromises genomic integrity. (A) Loss of MEF2A triggers the induction of DNA damage markers. AC16 cells were transfected with either NC or *MEF2A* target dsiRNA. In parallel, cells were stimulated with the topoisomerase I inhibitor, Etoposide, or the SERCA inhibitor, thapsigargin. Whole cell lysates were harvested 24 h post treatment and MEF2A, γH2AX, and Actin protein expression were measured by western blot. Images are representative of 3 independent experiments. (B) Representative micrographs of micronuclei detected by confocal microscopy of DAPI stained DNA in AC16 cells transfected with NC or *MEF2A*-targeting dsiRNA, or cells treated with nocodazole as positive control for micronuclei formation. (C) Quantification of micronuclei formation following MEF2A depletion. Bar graphs represent the percentage of micronuclei positive cells following NC (grey), *MEF2A* KD (red) or treatment with nocodazole (orange). Data is representative of 4 independent experiments and error bars represent SEM. **** p ≤ 0.0001 and ** p ≤ 0.01 as determined by one-way ANOVA. (D) Validation of cGAS knockout by assessment of 2’,3’ cGAMP production. Wild-type (H1) and cGAS KO cell lines were transfected with calf-thymus DNA (green bars, CT-DNA) or mock transfected (grey bars). Bar graphs represent average synthesis of 2’,3’ cGAMP as measured by ELISA across three independent experiments and error bars represent SEM. (E) The expression of cGAS is dispensable for the induction of inflammation following MEF2A loss. AC16 expressing a non-targeting sgRNA (H1) or clones of cells in which *CGAS* has been targeted (cGAS KO) were transfected with NC or *MEF2A* targeting dsiRNA. 24 h post transfection whole cell lysates were assessed for expression of phosphorylated STAT1 (Y701), total STAT1, cGAS, IRF1, MEF2A and GAPDH. Western blots are representative of 3 independent experiments. (F) cGAMP synthesis following *MEF2A* KD. AC16 cells were transfected with NC (black) or *MEF2A*-targeting dsiRNA (red) for 24 h. Bar graphs represent average synthesis of 2’,3’-cGAMP as measured by ELISA across 3 independent experiments. Error bars represent SEM. (G) AC16 H1 (clone1) and *IFI16* KO (clone 18) cells were transfected with NC or *MEF2A* targeting siRNA. 24 h post transfection relative expression of *IFNB1* mRNA was quantified by qPCR. Bar graphs represent *IFNB1* mRNA expression relative to *HPRT1* and H1 siMEF2A cells (value 100). Each data point represents values across 2 independent experiments and error bars represent SEM.

The accrual of DNA breaks can result in an accumulation of extranuclear DNA in the form of micronuclei. Experimentally, inhibition of microtubule dynamics by treatment with nocodazole promotes the mis-segregation of chromosomes (**Figure 4B**). We assessed whether the DNA damage response associated with MEF2A depletion was accompanied by accumulation of chromosomal aberrations by quantifying the percentage of cells with noticeable micronuclei. Compared with non-targeted controls, depletion of MEF2A led to a significant increase in the percentage of cells containing micronuclei, as was also observed for cells treated with nocodazole (**Figure 4C**). Thus, loss of MEF2A compromises genomic integrity, promoting cellular stress via the STING-IFN axis.

The accumulation of cytosolic nucleic acid or the loss of nuclear DNA compartmentalization is sensed by cGAS, and results in the synthesis of 2’,3’-cGAMP. To assess if cGAS was required for IFN induction upon *MEF2A* silencing, we generated *CGAS* KO cells by CRISPR-Cas9 genome editing (**Figure 4D and E**). Functional ablation of cGAS was confirmed by their lack of 2’,3’-cGAMP synthesis upon calf-thymus DNA transfection (**Figure 4D**). We observed that cells lacking cGAS expression were impaired in 2’,3’-cGAMP synthesis, while there was detectable accumulation of cGAMP in WT (H1) non-targeted control cells (**Figure 4D**). We then silenced MEF2A in either H1 or *CGAS* KO cells and examined STAT1 phosphorylation as a readout for STING activation. Robust levels of STAT1 Y701 phosphorylation upon MEF2A depletion were unaffected by cGAS deletion. The cGAS mutant cells also expressed comparable levels of IRF1, a type I IFN inducible protein (Forero et al., 2019) (**Figure 4E**). Importantly, relative to controls, production of 2’,3’-cGAMP by the cGAS mutant cells was unaffected after *MEF2A* silencing, suggesting that STING activation was driven by a cGAS and cGAMP-independent mechanism (**Figure 4F**). Because IFI16 can mediate non-canonical induction of type I IFN responses in a cGAS-independent, STING-dependent manner (Dunphy et al., 2018), we generated IFI16 KO cells (**Figure S4B**). Similar to data from cGAS-deficient counterparts, IFI16 KO and control cells induced comparable levels of *IFNB1* mRNA upon MEF2A depletion (**Figure 4G**). We conclude that MEF2A expression is necessary to prevent accumulation of DNA damage responses in the steady-state, which would drive inflammation using a STING-dependent, but cGAS- and IFI16-independent pathway.

### DDX41 detects R-loops that accumulate upon MEF2A depletion

DDX41 binds dsDNA and RNA:DNA hybrids and activates IFN (Stavrou et al., 2018; Zhang et al., 2011). Accordingly, we generated DDX41-deficient cells by CRISPR-Cas9 genome editing and measured inflammatory responses upon MEF2A depletion. IRF3 activation or STAT1 phosphorylation were attenuated dramatically upon silencing of MEF2A in DDX41-defficient cells (**Figure 5A**). Consistent with these findings, *DDX41* KO cells showed decreased STING phosphorylation after *MEF2A* knockdown (**Figure 5B**) relative to H1 controls. Decreased activation of STING, IRF3 and STAT1 was accompanied by a lack of *IFNB1* mRNA following *MEF2A* silencing in DDX41 KO cells (**Figure 5C**).

**Figure 5.**
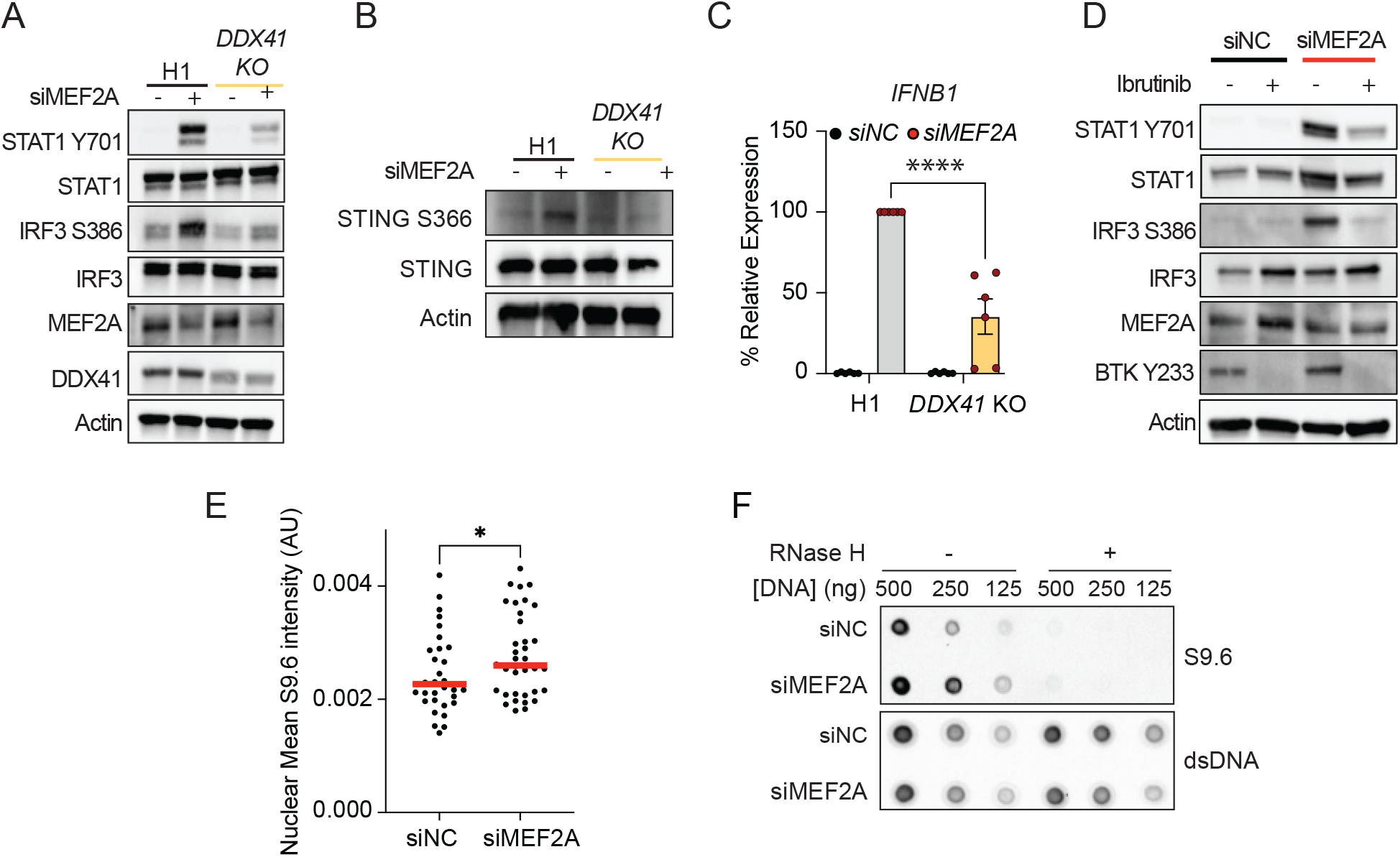
MEF2A triggers R-loop accumulation and DDX41-mediated inflammation-. (A) DDX41 is required for the induction of IFN responses following *MEF2A* depletion. AC16 *DDX41-* deficient cells were generated by CRISPR-Cas9 genome editing (DDX41 KO). Wild type (H1) and *DDX41* KO cells were transfected with NC or *MEF2A*-targeting dsiRNA for 24 h post transfection. Whole cell lysates were probed for protein expression of phosphorylated STAT1 (Y701), total STAT1, IRF3 (S386), total IRF3, MEF2A, DDX41 and Actin. Western blots are representative of a minimum of 3 independent experiments. (B) DDX41 is required for STING activation. Control cells (H1) and DDX41 KO cells were transfected with NC or *MEF2A-*targeting dsiRNA as indicated above. Phosphorylation of STING (S366) phosphorylation, total STING, DDX41 and Actin protein expression were probed by Western blot. Images are representative of 3 independent experiments. (C) AC16 H1 and *DDX41* KO cells were transfected with NC or *MEF2A*-targeting dsiRNA for 24 h prior to harvesting total RNA. Percent relative expression of *IFNB1* mRNA was quantified by qPCR. Bar graphs represent percent *IFNB1* mRNA expression relative to *HPRT1* control and normalization to *IFNB1* mRNA expression in H1 cells transfected with *dsiMEF2A* (100 %). Bar graphs represent average expression across 5 independent experiments and error bars represent SEM. **** p ≤ 0.0001 as determined by two-way ANOVA. (D) Inhibition of BTK abrogates the induction of IFN responses in U937 cells. U937 cells were treated with 4μM ibrutinib, specific kinase inhibitor for BTK, 2 h prior to dsiRNA transfection. Whole cell lysates were prepared 24 h post NC or *MEF2A* targeting and the expression of phosphorylated STAT1 (Y701), total STAT1, IRF3 (S386), total IRF3, MEF2A, phosphorylated BTK (Y233) and Actin. Image is representative of 3 independent experiments. (E) MEF2A loss leads to the accumulation of nuclear R-loops. AC16 cells were transfected with NC or MEF2A targeting dsiRNA for 24 h, cells treated with RNase T1 and RNase III and subjected to immunostaining with S9.6 antibody that recognizes RNA:DNA hybrids. Nuclear S9.6 foci were measured by confocal microscopy. Graph represents the average fluorescence intensity of nuclear foci across 30 nuclei depicted by each dot. * Represents p < 0.05 as determined by students t-test. (F) In vitro assessment of RNA:DNA hybrid formation after *MEF2A* knockdown. AC16 cells were transfected with NC or *MEF2A* targeting dsiRNA prior to extraction of genomic DNA. Serial dilutions of RNAse III treated DNA were dotted and cross-linked onto a nitrocellulose membrane as indicated. Membranes were incubated with S9.6 antibody to measure R-loops or dsDNA-specific antibodies as loading control. In addition, samples were mock-treated or digested with RNaseH to promote R-loop degradation.

An array of post-translational DDX41 modifications contribute to its nucleic acid binding capacity, including phosphorylation at tyrosine 414 (Y414) by Bruton’s tyrosine kinase (BTK) (Lee et al., 2015). Having demonstrated the genetic requirement for DDX41 in STING activation following MEF2A depletion, we asked whether chemical inhibition of BTK could abrogate this inflammatory response in U937 cells. Differentiation of U937 cells into monocytes by PMA treatment can promote basal BTK activity (**Figure S5A**). Like cardiomyocytes, depletion of MEF2A in differentiated U937 cells induces IRF3 and STAT phosphorylation (**Figure S5A**). In addition, loss of MEF2A expression in U937 promotes IFN secretion, as determined by the ISGF3 reporter assay (**Figure S5B**). Treatment of U937 cells with the BTK-specific kinase inhibitor, Ibrutinib, followed by knockdown of MEF2A significantly reduced phosphorylation of IRF3 and STAT1 relative to vehicle-treated cells in which MEF2A was depleted (**Figure 5D**). Thus, DDX41 and its activation are required for the induction of STING-mediated inflammation in the context of MEF2A silencing.

Recent studies have suggested that DDX41 is recruited to R-loops (Mosler et al., 2021; Nguyen et al., 2017; Weinreb et al., 2022), nucleic acid structures that render the genome susceptible to DNA breaks (Crossley et al., 2019). To determine whether MEF2A loss could promote R-loops accumulation, we conducted immunofluorescence-based assessment of R-loop accumulation in cells after knockdown of MEF2A, using an antibody specific for RNA:DNA hybrids (S9.6). Importantly, all cells were treated with RNase T1 and RNase III to minimize antibody cross reactivity with single-stranded and double-stranded RNA respectively (Smolka et al., 2021). Overall, we observed a significant increase in the intensity of nuclear S9.6 foci in *MEF2A*-depleted cells relative to non-targeted control cells (**Figure 5E**). Using an orthogonal approach, we measured the accumulation of R-loops *in vitro* by assessing the cross reactivity of S9.6 antibodies with genomic DNA isolated from *MEF2A*-depleted cells compared to controls (**Figure 5F**). Again, we observed an enhanced level of S9.6 antibody reactivity in nucleic acid samples derived from the MEF2A depleted cells. Importantly, treatment with RNase H, which specifically degrades RNA:DNA hybrid strands, resulted in the near complete loss of S9.6 staining, further confirming an accumulation of R-loops in muscle cells lacking MEF2A. As an additional control, neither silencing of MEF2A nor treatment with RNAseH affected dsDNA-specific antibody reactivity in any of the samples. Taken together, our data indicate that RNA:DNA hybrids are sensed in MEF2A depleted cells by DDX41, which promotes STING-mediated inflammatory responses.

### ATR kinase is pivotal for production of IFN following MEF2A depletion

The accumulation of R-loops activates DNA damage responses (DDR) coordinated by the kinases ATM and ATR, which ultimately promotes R-loop resolution (Chen et al., 2018; Hodroj et al., 2017; Kabeche et al., 2018; Matos et al., 2020). In addition, the DDR kinases have been implicated in the regulation of IFN following genotoxic stress (Burleigh et al., 2020; Dunphy et al., 2018; Forero et al., 2014). Thus, we addressed whether DDR kinase activity was necessary for the induction of inflammation in response to cellular stress elicited by decreased MEF2A expression. We pre-treated human fibroblasts with selective inhibitors of ATR (ATRi, ETP46464), ATM (ATMi, KU-55933) or DNA-PKcs (DNA-PKi, NU7441) kinase activity two hours prior to MEF2A depletion. Inhibition of ATR kinase activity specifically muted the activation of both IRF3 and STAT1 in response to the loss of MEF2A (**Figure 6A**). In contrast, robust phosphorylation of IRF3 and downstream STAT1 activation were observed when cells were treated with ATM kinase inhibitors as previously reported (Härtlova et al., 2015). Inhibition of DNA-PK kinase activity led to a minimal increase in basal IRF3 and STAT1 phosphorylation but did not affect STAT1 activation in response to MEF2A knockdown (**Figure 6A**). In addition, we observed that decreased MEF2A expression enhanced phosphorylation of the ATR substrate, RPA32, at serine 33 phosphorylation (Maréchal and Zou, 2013), and pre-treatment of cells with a highly specific ATR kinase inhibitor (AZD6738) led to a marked decrease in basal RPA32 phosphorylation. AZD6738 pretreatment also prevented increases in RPA32 phosphorylation in response to MEF2A depletion (**Figure 6B**). Importantly, treatment of fibroblasts with AZD6738 decreased the relative induction of *IFNB1 mRNA* in MEF2A depleted cells (**Figure S6B**).

**Figure 6.**
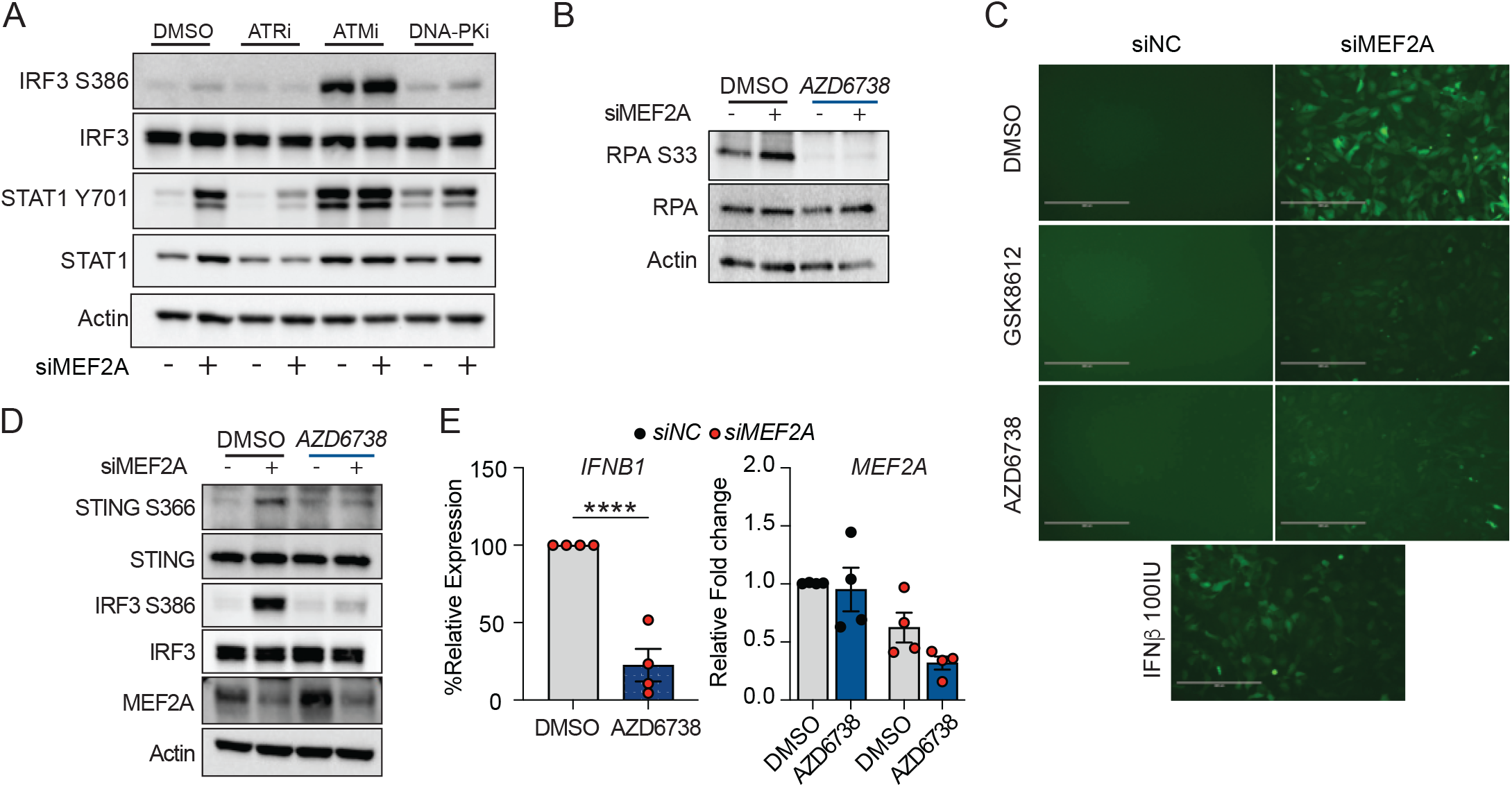
The inflammatory response to MEF2A loss required ATR kinase activity. (A) Contribution of specific DNA damage response kinase activity to the inflammatory response induced by *MEF2A* depletion. Human fibroblast cells, BJ/TERT, were pre-treated with ATR (ETP-46464, 2μM) ATM (KU-55933, 30μM), DNA-PK (NU7441, 4μM) kinase inhibitors or DMSO as vehicle control for 2 h prior to transfection with NC or *MEF2A*-targeting dsiRNA. Whole cell lysates were harvested 24 h post transfection and expression of phosphorylated IRF3 (S386), total IRF3, phosphorylated STAT1 (Y701), total STAT1, and Actin were measured by western blot. Data is representative of 3 independent experiments. (B) Phosphorylation of ATR substrate RPA32 following *MEF2A* depletion and DDR kinase activity inhibition. BJ/TERT cells were pre-treated with AZD6738 (2μM) for 2 h prior to transfection with dsiRNA. Whole cell lysates were probed for phosphorylated RPA32, total RPA32, and Actin by western blot. (C) Inhibition of TBK1 or ATR kinase activity impairs the induction of IFN responses in AC16 cells. AC16 ISRE-GFP reporter cell lines were pre-treated with TBK1 (GSK8621; 10μM) or ATR (AZD6738; 2μM) kinase inhibitors for 2 h prior to transfection with NC or *MEF2A*-targeting dsiRNA. As control, cells were stimulated with 100 IU/ml of recombinant IFNβ. Fluorescent signal was detected by epifluorescent microscopy. (D) The activation of STING in response to MEF2A loss requires ATR kinase activity. AC16 cells were pre-treated with ATR (AZD6783; 2μM) kinase inhibitors for 2 h prior to transfection with NC or *MEF2A*-targeting dsiRNA for 24 h. Whole cell lysates were harvested and expression of phosphorylated STING (S366), total STING, phosphorylated IRF3 (S386), total IRF3, MEF2A, and Actin were measured by western blot. Data is representative of 3 independent experiments. (E) ATR kinase inhibition impairs IFN production in response to MEF2A silencing. Bar graphs represented the percent relative expression of endogenous *IFNB1* mRNA expression in AC16 cells transfected with NC or *MEF2A*-targeting dsiRNA in the response to ATR kinase inhibition as described above (left). Relative *IFNB1* mRNA expression was calculated relative to expression in DMSO (value 100%) treated cells and normalized to *HPRT1* control. Relative *MEF2A* expression was normalized to *HPRT1* expression and DMSO treated cells (value 1). Each data point represents values across 4 individual experiments and error bars represent SEM. **** p ≤ 0.0001 as determined by student’s t-test.

We further defined the requirement for ATR kinase activity in IFN induction in cardiac muscle cells by generating a fluorescent cell-based assay (AC16-ISRE-GFP) in which activation of ISRE-mediated transcription drives the expression of green fluorescent protein (GFP) (Froggatt et al., 2021). As a primary control, IFN treatment of AC16-ISRE-GFP cells resulted in an increased expression of GFP (**Figure S6C**). Importantly, depletion of *MEF2A* in these cells also promoted GFP expression (**Figure S6D, left**), but did not affect cellular viability relative to non-targeted cells (**Figure S6D, right**). We further examined whether ATR kinase activity was also required to sustain IFN responses, as observed in fibroblast cells. Cardiac muscle cells were either stimulated with recombinant type I IFN or pre-treated with AZD6738. As a control, we pre-treated AC16-ISRE16 cells with the TBK1 kinase inhibitor, GSK8612, prior to MEF2A silencing. Overall, we observed GFP accumulation following transfection of cells with MEF2A-targeting siRNA, and was comparable to that observed in cells treated with IFN (**Figure 6C**). Pre-treatment of cells with either TBK1 or ATR inhibitor muted the expression of GFP in response to the loss of MEF2A. Treatment with hydroxyurea (HU) can induce the activation of ATR (Bianchi et al., 1986). Stimulation of cells with low dose HU induced ISRE activity (**Figure S6E**) and STING phosphorylation in an ATR dependent manner (**Figure S6F**).

In this regard, we next assessed whether ATR inhibition regulated the activation of STING in the context of MEF2A-depletion. Consistent with the results derived in human fibroblasts, treatment of AC16 cells with AZD6738 prior to siRNA transfection inhibited STING and IRF3 phosphorylation (**Figure 6D**). Finally, we determined whether the loss of STING and IRF3 activation following ATR inhibition led to decreases in *IFNB1* mRNA expression. Indeed, *IFNB1* mRNA induction was significantly reduced in AZD678 pre-treated cells relative to the levels detected in DMSO-treated cells (**Figure 6E**). Thus, ATR kinase activity positively regulates the STING-mediated inflammatory response to the loss of MEF2A. Collectively, we have shown that MEF2A is critical for maintaining genomic stability and prevents the activation of a discrete ATR-dependent DDR across cell types. This response feeds into the DDX41-STING pathway to promote inflammatory responses.

## DISCUSSION

In this study, we have identified MEF2A as a novel regulator of genomic stability that suppresses aberrant IFN activation. Downregulation of MEF2A expression resulted in the accumulation of R-loops, triple-stranded structures composed of a ssDNA and RNA:DNA hybrids. We also observed chromosomal breaks, as evidenced by micronuclei accumulation, increases in γH2A.X, and enhanced phosphorylation of the ATR substrate, RPA32. Importantly, inhibition of ATR kinase activity mitigated the induction of IFN responses upon MEF2A depletion. Our study also demonstrated that the accumulation of R-loops can induce IFN in a STING-dependent manner. Notably, we showed that the induction of STING required the expression and activation of the RNA:DNA sensor DDX41. Genetic ablation of *DDX41* or chemical inhibition of the DDX41 regulatory kinase, BTK, prevented the spontaneous production of IFN that follows the loss of MEF2A. These findings position the transcription factor MEF2A as an enforcer of genomic stability that, when compromised, leads to stress-induced, IFN-mediated inflammation. We found that the acute loss of MEF2A results in the induction an antiviral state across multiple cell types, including cardiomyocytes, fibroblasts, and monocytes.

Depletion of MEF2A expression induced both type I and type III IFN expression. Although these two IFN families promote an antiviral state in target cells, expression of type I and III IFN receptors varies across tissue types. The type I IFN receptor is expressed on all nucleated cells while, the type III IFN receptor expression is restricted primarily to epithelial barriers and a subset of immune cells (Dowling and Forero, 2022). We used orthogonal approaches to demonstrate that cardiac muscle cells respond primarily to type I IFNs. Treatment of cells with recombinant type III IFN failed to induce the expression of ISGs, while blockade of secreted type I IFN by treatment with B18R was sufficient to abrogate the induction of antiviral responses. The translational potential of our findings is underscored by emerging virus outbreaks, which have refocused the attention on the use of recombinant IFN therapies for the clinical management of viral disease (Beyer and Forero, 2022). In addition, induction of IFN responses during ischemic injury to cardiomyocytes can induce STING/IRF3-dependent inflammatory damage (Calcagno et al., 2020; King et al., 2017). Our findings suggests that type I IFN targeted therapies may have a greater benefit in the management of cardiotropic virus infections. Conversely, type III IFN induction might contribute less to adverse outcomes after myocardial infarction, compared with type I IFN.

MEF2 proteins are transcriptional regulators implicated in the control of inflammatory responses. In primary murine microglial cells, MEF2C expression is induced by type I IFN and it serves to dampen age related inflammatory responses (Deczkowska et al., 2017). We observe similar increase in *MEF2C* mRNA expression in cells with active IFN responses. In vitro, microglial expression of MEF2D is required to sustain *Irf7* expression (Lu et al., 2021). In murine macrophages, MEF2A is necessary to promote chromatin accessibility at the *Ifnb1* locus. Deletion of *Mef2a* suppresses the induction of *Ifnb1* following LPS stimulation (Cilenti et al., 2021). Whether MEF2 transcription factors regulate inflammatory responses in human cells was less understood. Our results using immortalized cardiomyocytes, fibroblast, and myeloid cells indicate that MEF2A serves indirectly as a negative regulator of inflammation by preventing cellular stress. Silencing of MEF2A led to the induction of DNA damage responses, as well as the accumulation of R-loops. Previous studies have demonstrated that murine MEF2A silencing can alter the expression profile of cell cycle regulatory genes, impacting the DNA content profile (Desjardins and Naya, 2017). Indeed, we observed a downregulation in mitotic regulatory genes upon transient depletion of these genes in human cardiac muscle cells. Importantly, our study correlates the inductions of cellular stress and IFN-mediated inflammation. Future studies should address how disruption of cell cycle progression by MEF2A depletion specifically promotes R-loops. Understanding the context in which nucleic acid mis-localization and lesion accumulation drives PRR activation is critical for the design and implementation of STING immunomodulatory therapies for the management of autoinflammation and cancer (Decout et al., 2021).

R-loops function to promote mitochondrial DNA replication, control transcriptional rates, coordinate faithful chromosome segregation, and facilitate immunoglobulin class switch recombination (Aguilera and Gómez-González, 2017; Kabeche et al., 2018; Yu et al., 2003). However, aberrant accumulation of R-loops can compromise chromosomal stability by rendering the genome susceptible to DNA breaks. Pathological R-loop accumulation is the hallmark of inflammatory neurodegenerative diseases such as amyotrophic lateral sclerosis (ALS), Fragile X syndrome, and AGS (Cristini et al., 2022; Park et al., 2021). Excessive R-loop formation is also observed in several proliferative malignancies. Cellular nucleases and helicases metabolize RNA:DNA hybrids to mitigate the cytotoxicity of their aberrant accumulation. The expression of Senataxin (SETX) and the related DEAxQ-like domain containing helicase Aquarius (AQR) (Nguyen et al., 2017; Sollier et al., 2014) is necessary to curb R-loop accumulation. DEAD/DEAH-box helicases, such as DDX5, DDX19, and DDX41 amongst others, also are recruited to R-loops and metabolize RNA:DNA hybrids (Cargill et al., 2021). On the other hand, DexH-box helicases, such as DHX9, both promote physiological R-loop formation at centromeres and drive their pathogenic accumulation during defective splicing (Chakraborty et al., 2018). Given the physiological importance of these helicases in the formation and resolution of R-loops, understanding the factors that control their recruitment, activation, and downstream effector functions should be further explored.

Proximity ligation studies show DDX41 occupancy at R-loops is necessary for their resolution (Mosler et al., 2021; Weinreb et al., 2021). Transgenic zebrafish models show that *ddx41* loss of function (LOF) mutations drive R-loop accumulation that can trigger cGAS-STING pathway dependent activation of inflammation (Weinreb et al., 2021, 2022). In this model, *ddx41* LOF disrupts splicing of cell cycle genes and can drive the activation of ATM and ATR coordinated DDR that inhibit erythropoiesis. In contrast, we found that in the context of acute MEF2A depletion, neither cGAS nor IFI16 appeared to coordinate the induction of downstream inflammatory responses. We found that in addition to these roles, the accumulation of R-loops induces DDX41-dependent activation of STING and type I IFN-mediated inflammation. However, DDX41 depletion did not result in a complete abrogation of *IFNB1* mRNA expression. It is possible that the accumulation of DNA breaks in DDX41-deficient cardiac muscle cells could trigger residual DDX41 in our cells or a cGAS/IFI16-dependent inflammatory pathway.

Consistent with our studies, accumulation of RNA:DNA hybrids triggered by MEF2A silencing also required ATR activation (Matos et al., 2020; Mosler et al., 2021). The activation of ATR by R-loop accumulation has been well established. Enhanced R-loop accumulation and replication fork reversal by MUS81 required ATR activation in cells lacking the splicing factor SRSF1. Similarly, the disruption of DDX41 expression can enhance R-loop abundance and subsequent activation of ATR (Mosler et al., 2021). Leveraging the MEF2A-depletion model, we showed that ATR kinase activity is necessary to coordinate the induction of inflammatory responses upon increased R-loop abundance. In contrast, impairment of other DDR kinases, ATM and DNA-PKcs kinase, had no significant impact on the induction of STING activation after MEF2A downregulation. ATR localizes to R-loops during mitosis to promote faithful chromosome segregation (Kabeche et al., 2018), and activation of the ATR/Chk1 pathway upon replication stress is important for the recruitment of nucleases into R-loops (Hodroj et al., 2017). A better understanding of the context in which ATR is recruited to R-loops allow us to define whether it has distinct roles in the activation and recruitment of RNA:DNA nucleases, such as DDX41, that either promote R-loop or induce inflammation.

In conclusion, we found that MEF2A regulates genomic stability to protect cells from unscheduled production of type I IFN. This study bridges the activation of the RSR upon loss of MEF2A with DDX41-dependent activation of STING-mediated inflammation. Specifically, we demonstrate that ATR kinase activity is necessary for the activation of STING activation following R-loop accumulation. As such, our findings reveal a novel role for MEF2A in the regulation of cellular homeostasis, and suggest a potential benefit for the use of specific ATR kinase inhibitors to mitigate deleterious inflammation in neurodegenerative and proliferative diseases defined by increases in R-loop abundance.

## MATERIALS AND METHODS

### Cell lines, cell culture conditions and treatments

Human cardiomyocyte (AC16) and derivate cell lines were grown in Dulbecco’s modified Eagle’s medium (DMEM) supplemented with 12.5% FBS, 2mM Glutamine, 100 U/ml Penicillin and 100 mg/ml Streptomycin and maintained at 37°C in 5% CO_2_. Non-targeted (H1), IRF3, STING, cGAS, IFI16 and DDX41-deficient AC16 cells were generated by CRISPR-Cas9 genome editing as previously described (Forero et al., 2019). Transduced cells were enriched using antibiotic selection. BJ/TERT human fibroblasts were cultured in DMEM supplemented with 10% FBS, 2mM glutamine, 100 U/ml penicillin and 100 mg/ml streptomycin and maintained at 37°C in 5% CO_2_. Huh7 human hepatoma cells and derivatives cell lines were cultured in DMEM supplemented with 10% FBS, 2mM Glutamine, 100 U/ml Penicillin and 100 mg/ml Streptomycin and maintained at 37°C in 5% CO_2_. U937 monocytes were cultured in RPMI 1640 media supplemented with 10% FBS, 2mM glutamine, 100 U/ml penicillin and 100 mg/ml streptomycin and maintained at 37°C in 5% CO_2_. THP-1 monocytes were maintained in RPMI 1640 media supplemented with 10% FBS, 2mM glutamine, 100 U/ml penicillin, 100 mg/ml streptomycin, 1 mM sodium pyruvate, 10 mM HEPES and 0.05 mM 2-mercaptoethanol and maintained at 37°C in 5% CO_2_. U937 and THP-1 cells were differentiated for 48 hours (h) in their respective complete media containing 40 nM phorbol 12-myristate 13-acetate (PMA) followed by resting for 24 h in RPMI 1640 supplemented with 1% FBS.

Cells were stimulated with recombinant IFNβ (PBL Assay Science) and IFNλ3 (R&D Systems) at the indicated concentrations. Specific kinase inhibitors targeting, ATR (ETP-46465, AZD6738) (Cayman Chemicals), ATM (KU-55933) (SelleckChem), DNA-PK (NU7441) (SelleckChem), and TBK1 (GSK8612) were used at the indicated concentrations (Supplementary Table 2). Recombinant B18R (Life Technologies) was used at 1ug/ml. Etoposide and thapsigargin (Millipore Sigma) as indicated.

### Plasmids and Oligonucleotides

CRISPR-Cas9 plasmids; pRRL-H1-PURO (non-targeting), pRRL-STING-PURO, pRRL-cGAS-PURO, pRRL-IRF3-PURO, pRRL-IFI16-PURO were a gift from Dr. Daniel Stetson (University of Washington) (Gray et al., 2016). The ISRE reporter plasmid, pISRE-sfGFP, was a gift from Dr. Nicholas Heaton (Duke University) and has been previously described (Froggatt et al., 2021). pRRL-DDX41-PURO was generated by cloning single-guide RNA (sgRNA) targeting DDX41 (5’-CCTCATCTTCCGCCTCGGAG-3’) into empty pRRL-Cas9-PURO plasmids as previously described (Forero et al., 2019; Gray et al., 2016). Gene silencing was conducted using dicer-substrate interfering RNA (dsiRNA) specific to MEF2A, MEF2C, MEF2D, TMEM173, MAVS, IRF1, STAT1 or non-targeting control (Supplementary Table 2). Transfection were carried out using 20 nM of dsiRNAs delivered intracellularly using TransIT-TKO (Mirus) according to manufacturer’s guidelines. Additional MEF2A targeting was done using silencer siRNA (Dharmacon) targeting MEF2A or scramble control (Supplementary Table 2).

### Viral infections and quantification

VSV-GFP stock was a gift from Dr. Michael Gale Jr (University of Washington) (Fredericksen and Gale, 2006). Sendai Virus (Cantell strain) was acquired from Charles River. AC16 cells were seeded in 24-well plates at a density of 2 x 10^5^ cells/well and transfected with negative control or MEF2A-targeting dsiRNA (IDT, Supplementary Table 2). Cells were infected with VSV-GFP as previously described (Forero et al., 2019). Following 24 h of infection, the culture medium was removed, cells were fixed with 4% PFA in PBS, and stained with Crystal violet stain (3% w/v) in 50% ethanol. Plates were imaged using the ChemiDoc XRS+ (BioRad) imaging system. Coxsackievirus B3 (CVB3)-Nancy was a kind gift from Dr. Raul Andino (University of California San Francisco) and was prepared as previously described (Laufman et al., 2019). AC16 cells were seeded in 12-well plates at a density of 3.5 x 10^5^ cells/well and transfected with negative control or MEF2A-targeting dsiRNA. Cells were infected in a minimal volume for 1h at 37°C in 5% CO_2_ followed by washing with PBS and culturing in full serum media for 24 h. CVB3 RNA quantification was conducted via SYBR green based qPCR using primers specific to CVB3 VP1 (Supplementary Table 2).

### RNA extraction and quantification of gene expression

Total RNA was extracted using the NucleoSpin RNA extraction kit (Macherey-Nagel) as indicated by manufacturer guidelines. cDNA synthesis was performed using the QuantiTect RT kit (QIAGEN) or iScript cDNA synthesis kit (BioRad) according to the manufacturer guidelines. Relative quantification of mRNA was done by qPCR using the ViiA7 qPCR system with TaqMan reagents (Life Technologies) or CFX-384 with SSO Advanced Probes reagents (BioRad) using the *HPRT1* as reference gene. Primers and probes used for qPCR assays in this study were acquired from IDT or Life Technologies as indicated in Supplementary Table 2.

### Western blot analysis

Whole cell lysates were prepared from cells using RIPA buffer (10 mM Tris-Cl (pH 8.0), 1 mM EDTA, 0.5 mM EGTA, 1% Triton X-100, 0.1% sodium deoxycholate, 0.1% SDS, 140 mM NaCl) supplemented with Halt protease and phosphatase inhibitor cocktail (Pierce). Protein quantification and normalization was done using the BCA Protein Assay Kit (Pierce). 10-30 ug total protein were resolved by SDS-PAGE and transferred to PVDF membranes (Bio-Rad). Primary antibody incubations were done overnight with antibodies diluted in 3% BSA in TBS-T (Tris-buffered saline/Tween 20), and species-specific HRP conjugated secondary antibodies. Chemiluminescent image acquisition was performed using a ChemiDoc Touch (BioRad).

### RNA sequencing, data processing, and analysis

AC16 cells were seeded in 24-well plates at a density of 2 x 10^5^ cells/well and transfected with either negative control or MEF2A-targeting dsiRNA in triplicate. Total RNA was extracted 24 h post transfection as described above. Total RNA fluorometric quantification of was done using the Qubit RNA BR assay kit (Invitrogen) and assessment of RNA integrity was performed using the RNA 6000 Nano Kit in the 2100 Bioanalyzer (Agilent). Library preparation, QC, and sequencing was carried out by Seattle Genomics (http://www.seattlegenomics.com). Briefly, the synthesis of cDNA libraries was conducted using the TruSeq Stranded mRNA Library Prep Kit. Libraries were sequenced using the Ilumina NextSeq 500 sequencer. Human genome sequence (fasta) and gene transfer files (gtf) were obtained through iGenomes (https://support.illumina.com/sequencing/sequencing_software/igenome.html). Raw RNaseq data (Fastq files) were demultiplexed prior to read quality determination (FastQC, version 0.11.3). Remaining ribosomal RNA reads were digitally removed using Bowtie2 (version 2.3.4). All samples had a minimum of twenty million reads per sample which were aligned to the human genome (GRCh37) using STAR (version 2.5.3a) and gene counts were derived using HTSeq (version 0.6.1). Gene counts were filtered (mean of ≥10 or greater across all samples) prior to statistical analysis using R statistical programming language (version 3.4.3) and ‘edgeR’ (version 3.20.9). Gene counts normalization was performed with voom and differential expression analysis was carried out with ‘limma’ (version 3.34.8). Gene Ontology analysis of differentially expressed transcripts (lfc |0.26|; p-value < 0.01) was performed using EnrichR.

### Micronuclei quantification assay

3 x 10^6^ AC16 cells were plated in a 100-mm dish and treated with 100 ng/mL of nocodazole for 6h. Mitotic cells were harvested by shake-off and washed 3x with PBS. Mitotic cells were then counted and plated in on poly-L-ornithine (PLO)-coated #1.5 12 mm glass-coverslips (Thomas Scientific). Simultaneously, untreated AC16s were plated on PLO-coated coverslips and transfected with 40 nM of non-targeting dsiRNA or dsiRNA targeting *MEF2A* for 24 h. Cells were fixed with ice-cold 100% methanol for 15 min at −20°C. Fixed cells were then washed 2 times with PBS and blocked in PBS containing 3% BSA and 0.3% Triton-X100. Cells were stained using Lamin B1 antibodies (CST) at 1:200 dilution for 1h at room temperature in PBS containing 1% BSA and 0.1% Triton X-100. Cells were washed 3 times and stained with Alexa-488 conjugated anti-rabbit secondary (Invitrogen) and DAPI (Thermo Scientific) for 1h at room temperature (RT) in PBS containing 1% BSA and 0.1% Triton X-100. Cells were washed and mounted with ProLong Glass Antifade mounting medium (Thermo Fisher). Samples were imaged using a Nikon Eclipse Ti laser scanning confocal microscope, 60x oil-immersion lens. Percent micronuclei was quantified by counting the number of micronuclei per nucleus in the field of view using FIJI (Schindelin et al., 2012).

### *In vivo* and *in vitro* R-loop quantification assay

RNA:DNA hybrid microscopy was conducted as previously described with some modifications (Kim et al., 2018). AC16 cells were seeded at 0.5 x 105 cells/well on poly-L-ornithine (PLO)-coated #1.5 12 mm glass-coverslips (Thomas Scientific) and transfected with 40 nM of siNC or siMEF2A for 24 h. Cells were fixed with ice-cold 100% methanol for 15 min at −20°C. Fixed cells were then washed 2 times with PBS and simultaneously blocked and subjected to RNase digestion for 2h at 37°C in PBS containing 3% BSA, 3mM MgCl2, 2U RNase T1 and 2U RNase III. Cells were then immunostained using S9.6 antibody (Kerafast) at 1:500 dilution for 1h at RT in PBS containing 1% BSA and 0.1% Triton X-100. Cells were washed 3 times and stained with Alexa-488 conjugated anti-mouse secondary (Invitrogen) and DAPI (Thermo Scientific) for 1h at room temperature in PBS containing 1% BSA and 0.1% Triton X-100. Cells were washed and mounted with ProLong Glass Antifade Mountant (Thermo Fisher). Samples were imaged using a Nikon Eclipse Ti laser scanning confocal microscope, 60x oil-immersion lens. Images were quantified using CellProfiler (Stirling et al., 2021) and presented as mean S9.6 foci intensity per nuclei.

RNA:DNA hybrid dot blots were performed as previously described (Mosler et al., 2021). In brief, total *MEF2A*-targeted and control cell DNA was extracted using NucleoSpin Tissue DNA kit (Machery-Nagel). DNA was digested with RNase III (Thermo Fisher Scientific) 1U/μg of DNA for 2h at 37°C followed by heat inactivation at 65°C for 20 min. Samples were then split in half and digested with RNaseH (Thermo Fisher Scientific) at 37°C overnight. RNAse treated DNA was then blotted on nitrocellulose membranes and UV-crosslinked using 1200 μJ x 100 cm^2^. Membranes were blocked for 30 min in 3% BSA in TBS and 0.1% Tween-20. Membranes were probed overnight at 4°C with anti-RNA:DNA hybrid (S9.6) (Kerafast) at 1:10,000 dilution or dsDNA antibody (Abcam) at 1:1000 dilution. Membranes were incubated for 45 min with anti-mouse IgG HRP conjugated secondary antibodies for 45 min. Membranes were imaged using Bio-Rad Chemidoc.

### ISGF3 Gaussia luciferase and ISRE GFP reporter assays

To generate the ISGF3 Gaussia Luciferase (Gluc) reporter construct, pTRIPZ-5xISGF3-BS-hGLuc-PEST, 5 tandem ISGF3 consensus sequences (5’-CGAAGAAATGAAACT-3’) (Schmid et al., 2010) were cloned with hGLuc-MODC-PEST into a pTRIPZ lentiviral plasmid (Supplementary Table 2). Lentivirus encoding the reporter was packaged used to transduce human hepatoma Huh7 cells prior to single-cell cloning of reporter cells. To assess the presence of secreted IFN from dsiRNA transduced AC16 cells, cell supernatants from *MEF2*-depleted and non-targeting control were harvested 24 h post transfection and transferred onto 5xISGF3-GLuc Huh7 reporter cells. Reporter cells were then incubated at 37°C and 5% CO_2_ for 24 h prior to assessment of Gaussia luciferase secretion into the media. Sample supernatants were diluted 1:1 with Gaussia Luciferase glow assay substrate (Thermo Fisher Scientific) and luminescence measured using a Synergy HTX (BioTek). To generate ISRE-GFP reporter cell lines, AC16 were stably transduced with lentivirus pISRE-sfGFP and pools were treated with 100IU/ml of recombinant IFNβ or transfected with non-targeting control or *MEF2A*-targeting siRNA in the presence of the indicated inhibitors prior to epifluorescent imaging using an EVOS cell imaging system (Thermo Fisher Scientific). For flow cytometric assessment of fluorescence expression, cells were recovered by trypsinization, labeled with Fixable Viability Dye eFluor™ 780 (Invitrogen) per manufacturer’s guidelines and fixed with 4% methanol-free PFA. Fluorescence intensity was using a CANTO analyzer (BD). Flow cytometry data was analyzed using FlowJo (TreeStar).

### ELISA quantification of 2’,3’-cGAMP production

H1 control AC16 cardiomyocytes and *cGAS* KO AC16s were seeded in 12-well plates and transfected with 5 mg of CT-DNA using Transit X2 (Mirus) at a 2:1 ratio for 8 h prior to harvest. Wild-type AC16 cells were seeded in 12-well plates prior to depletion of *MEF2A* with dsiRNA or non-targeting control as described above for 24 h prior to harvest. 2’,3’-cyclic GMP-AMP (cGAMP) was quantified using the Direct 2’,3’-cGAMP ELISA Kit (Arbor Assays) according to manufactures protocol. Briefly, cells were lysed in 150ul of sample diluent solution for 15 min at room temperature. Cell lysates were recovered by scrapping and cell debris was removed by centrifugation at 700 x g at 4°C. Absorbance was read at 450 nm using a Synergy HTX (BioTek).

### Quantification and Statistical Analysis

Statistical analysis was performed using GraphPad Prism 9.0 (GraphPad software La Jolla, CA). Statistical significance was calculated as indicated for each experiment and across all experiments, p-values of < 0.05 were considered significant and are indicated by asterisks (*).

### Public data availability

The data generated in this study are available via the following accession identifiers on the NCBI-GEO database (GEO: GSE209601).

## ACKNOWLEDGEMENTS

We thank Daniel B. Stetson, Joshua Woodward, and Haitao Wen for sharing reagents and helpful discussions. We thank Richard Robinson and Eugene Oltz for critical reading of the manuscript. This work was supported in part by T32HL007312, The Ohio State University institutional funds (A.F.), The Ohio State University Comprehensive Cancer Center and the National Institutes of Health under grant number P30 CA016058, and R21AI141823 (R.S.). The content is solely the responsibility of the authors and does not necessarily represent the views of the funding agencies.

## AUTHOR CONTRIBUTIONS

Investigation and Formal Analysis (J.R.S, J.W.D, A.K., J.S., and A.F.); Conceptualization (J.W.D, J.R.S, A.F.), Writing (J.R.S, A.F.), Supervision (R.S. and A.F.), Funding Acquisition (R.S. and A.F.).

