## Supplementary material for "MEF2A suppresses replicative stress responses that trigger DDX41-dependent IFN production": Data Supplement

**Supplemental Figure 1. Induction of IFN responses by MEF2A depletion across cell types.** (A) Validation of IFN induction by various *MEF2A* siRNA sequences. AC16 cells were transfected with siRNA (Dharmacon) or dicer-substrate siRNA (dsiRNA) targeting distinct regions of *MEF2A* for 24 h prior to harvesting whole cell lysates. Control cells were transfected with equimolar concentrations of non-targeting siRNA. In all cases, silencing of *MEF2A* led to an increase in STAT1 phosphorylation (Y701). (B) Expression of MEF2 transcription factor mRNA in human left ventricular cardiomyocytes. Heatmap represents the transcript per million detected by RNA sequencing as reported in GTEx. (C) Expression of MEF2 transcripts in human fibroblast cell line, BJ/TERT. Bar graphs represent average *MEF2A* (red), *MEF2C* (blue) and *MEF2D* (yellow) mRNA expression relative to *HPRT1* across 2 independent experiments. *MEF2B* expression was not detected in these cells (ND). (D) Depletion of *MEF2A* (green) promotes *IFNB1* (black) and *CXCL10* (red) in BJ/TERT cells relative to *HPRT1* and NC (left, value 1). Each data point represents an individual experiment, n=3. Depletion of *MEF2A* (green) promotes *IFNB1* (black) and *CXCL10* (red) in PMA-differentiated THP-1 monocytes relative to *HPRT1* and NC (right, value 1). Each data point represents an individual experiment, n=3.

**Supplemental Figure 2. MEF2A depletion promotes spontaneous IFN responses and sensitizes cells to exogenous type I IFN treatment.** (A) Validation of ISGF3 responsiveness of reporter cell line. Bar graph represents *Gaussia* luciferase activity detected from 5xISGF3-Gluc Huh7 cells stimulated with recombinant IFNβ (500 U/mL, pink) or IFNλ3 (500 ng/mL, green) or mock treated for 24 h. (B) AC16 cells were transfected with siRNA targeting MEF2A, STAT1, and IRF1 or NC. After 24 h of transfection, cells were stimulated with 25 IU/ml of recombinant IFNβ for 30 mins. Protein phosphorylation of IRF3 (S386) and STAT1 (Y701) was assessed by western blot.

**Supplemental Figure 3. *MEF2A* depletion induction of IFN requires STING but not MAVS.** (A) Induction of type III IFN and IFN-stimulated genes by DNA and RNA transfection in AC16 cells. AC16 cells were transfected with calf-thymus DNA (DNA, purple) or HCV PAMP (RNA, blue) for 6 h prior to total RNA harvest. Bar graphs represent average relative fold changes in mRNA levels were calculated relative to *HPRT1* and normalized to mock-transfected cells (value 1). Bar graph represents 3 independent experiments and error bars represent SEM. (B) Induction of IFN in response to exogenous dsRNA and virus infection. AC16 cells were treated with the indicated doses of floating poly(I:C) (TLR3 ligand) or Sendai Virus (Cantell strain). Bar graphs represent relative fold changes in mRNA levels were calculated relative to *HPRT1* and normalized to mock-transfected cells (value 1). (C) Confirmation of STING depletion following *TMEM173* targeting by CRISPR-Cas9 gene-editing technologies. H1 non-targeting control and *TMEM173* targeted cells were mock-treated or stimulated with IFNβ (25IU/ml) for 24 h. Total lysates were probed for STING and Actin protein expression by western blot. The numbers indicate clonal cell populations.

**Supplemental Figure 4. *Response to ER stress and IFI16 deletion in AC16 cells.*** (A) ER stress response elicited by treatment of AC16 cells with topoisomerase I (etoposide) and SERCA inhibitors (thapsigargin) or *MEF2A*-targeting. Cells were treated as described in Figure 4A. Whole cell lysates were probed for ATF4, spliced XBP1 (XBP1s), and Actin protein expression. Western blots representative of 3 independent experiments. (C) Generation of IFI16 knock-out cells. AC16 H1 and *IFI16* KO cells were generated by transduction with Cas9 and non-targeting sgRNA (H1) or IFI16-targeting sgRNA. Clonal lines were derived as indicated by individual numbers and expression of IFI16, STING, IRF3, MEF2A, STAT1 and Actin was assessed by western blot. Cone 1 H1 cells and clone 18 IFI16 KO cells were selected to assess response to *MEF2A* silencing.

**Supplemental Figure 5. Induction of inflammatory responses in macrophage cells upon *MEF2A* silencing.** (A) Bruton's tyrosine kinase (BTK) activity and MEF2A knockdown in U937 cells. Differentiation of U937 cells with 48 h PMA treatment (40 ng/ml) enhances BTK activity and STAT1 expression. PMA-differentiated cells were transfected with NC control or *MEF2A*-targeting dsiRNA for 24 h prior to harvesting whole cell lysates. Protein expression was assessed by western blot. (B) IFN secretion from *MEF2A* depleted U937 cells. Bar graphs represent average *Gaussia* luciferase activity detected from 5xISGF3-GLuc Huh7 cells stimulated with supernatants from U937 human monocyte cells transfected with either NC or *MEF2A* targeting siRNA. Each data point represents an individual experiment, n=2.

**Supplemental Figure 6. ATR is required for IFN induction following replicative stress.**
(A) Changes in endogenous *IFNB1* mRNA expression following ATR kinase inhibition in BJ/TERT cells transfected with NC or *MEF2A*-targeting siRNA. BJ/TERT cells were pre-treated with ATR kinase inhibitors (AZD6783; 2μM) for 2 h prior to 24 h transfection with NC or *MEF2A*-targeting siRNA. Bar graphs represent the average percent induction of *IFNB1* mRNA following *MEF2A* silencing in AZD6738 treated cells relative to DMSO (value 100%) or AZD6738 and expression level of *MEF2A* relative to DMSO (value 1) (left). Each data point represents values across 2 individual experiments and error bars represent SEM. (B) Characterization of AC16-ISRE-GFP

reporter cell lines. Cells were treated with 500IU of recombinant IFNβ for 24 h prior to detection of FITC expression by FACS. (C) Induction of ISRE activity by loss of MEF2A expression. AC16-ISRE-GFP cells were transfected with NC or MEF2A-targeting dsiRNA or 24 h prior to detection of FITC (left) or fluorescent-dye uptake (APC-Cy7) to assess cellular viability (right). (D) The induction of replicative stress by hydroxyurea treatment induces IFN responses. AC16-ISRE-GFP cells were stimulated with DMSO or 100nM hydroxyurea (HU) for 30 h. Fluorescent micrographs are representative of 3 independent experiments. (E) The phosphorylation of STING following hydroxyurea treatment requires ATR kinase activity. AC16-ISRE-GFP cells were stimulated with DMSO or 100nM hydroxyurea (HU) for 30 h in the presence or absence of ATR inhibitor (AZD6738; 2μM). Whole cell lysates were harvested and protein expression of phosphorylated CHK1, phosphorylated STING, total STING and Actin were assessed by western blot. Images are representative of 3 independent experiments.

**Supplemental Table 1. Differentially expressed genes after MEF2A silencing.** List of differentially expressed genes (LFC |0.26|; adj p-value 0.01) following knockdown (KD) of *MEF2A* with siRNA.

**Supplemental Table 2. Key reagents and resources**

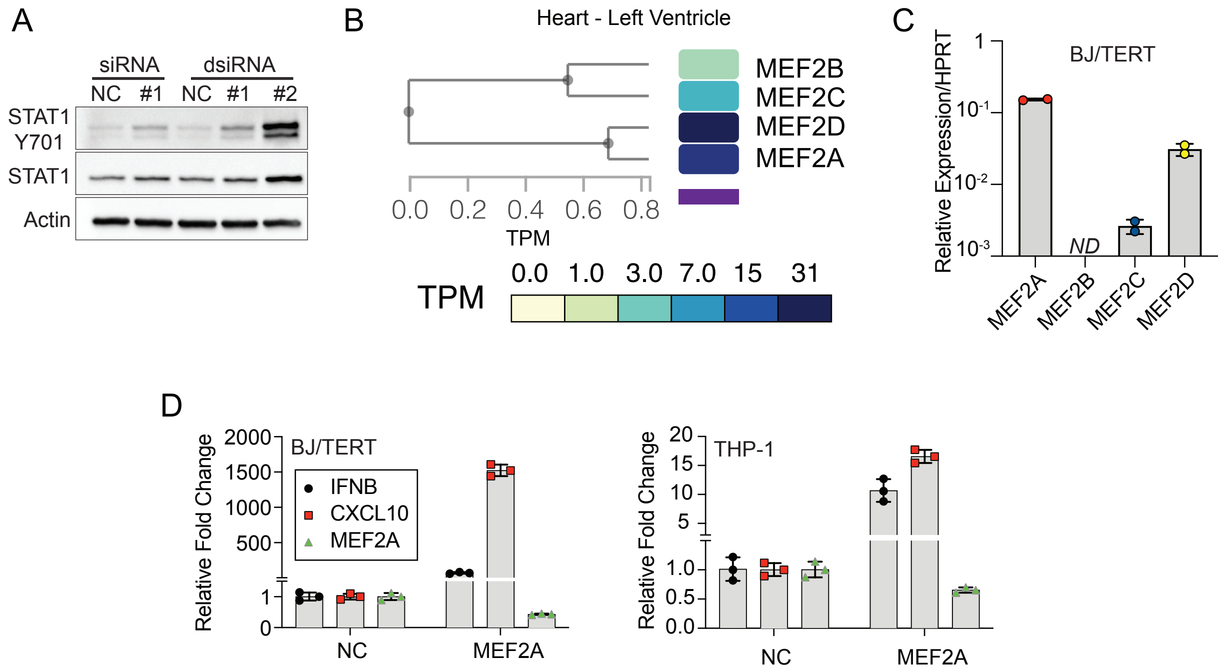


Supplementary Figure 1

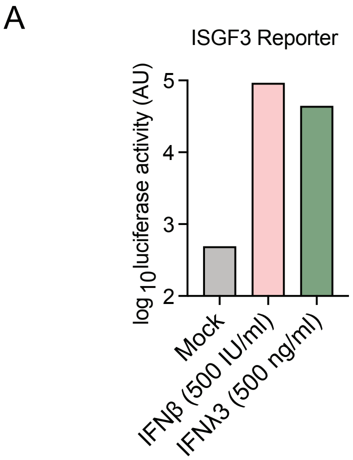


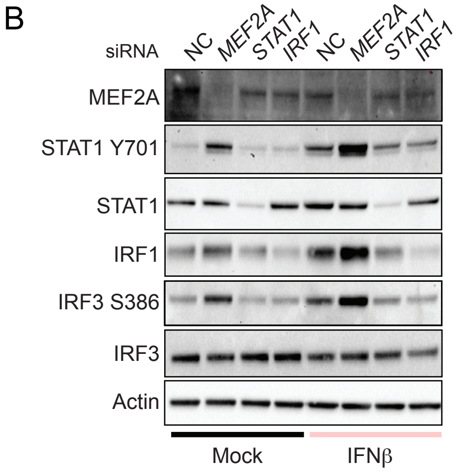


Supplementary Figure 2

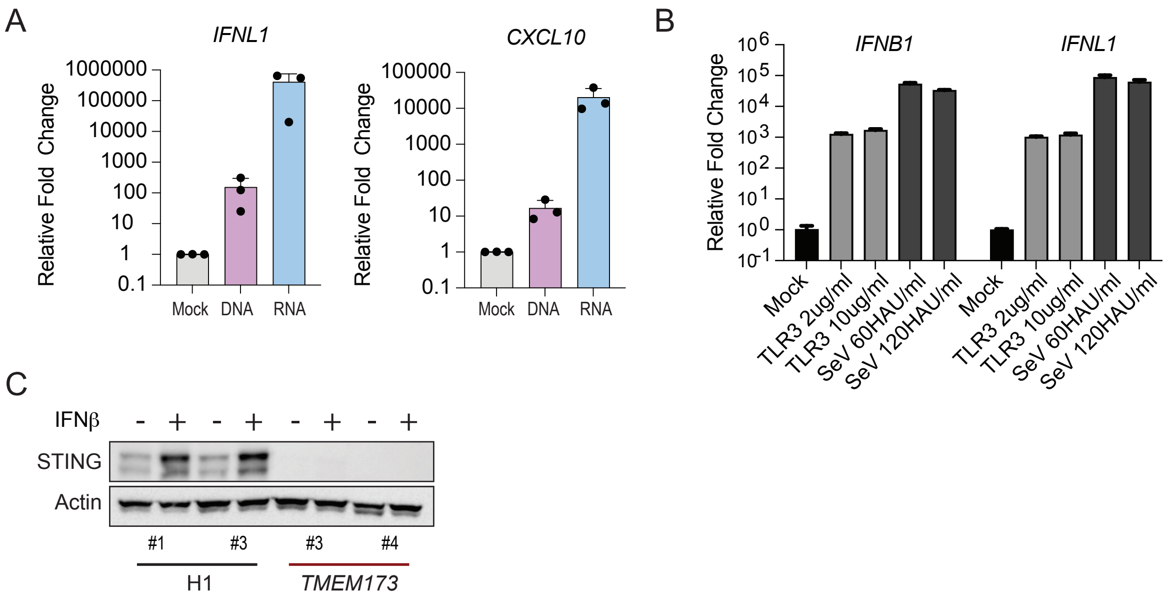


Supplementary Figure 3

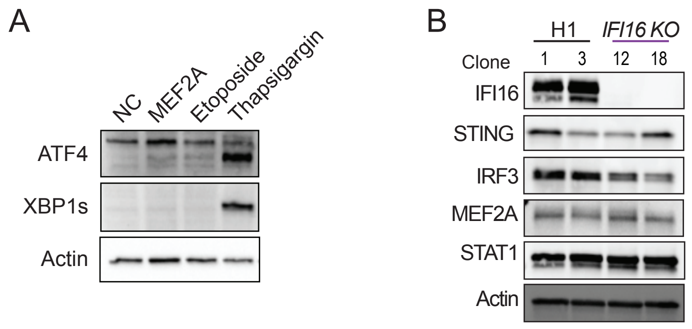


Supplementary Figure 4

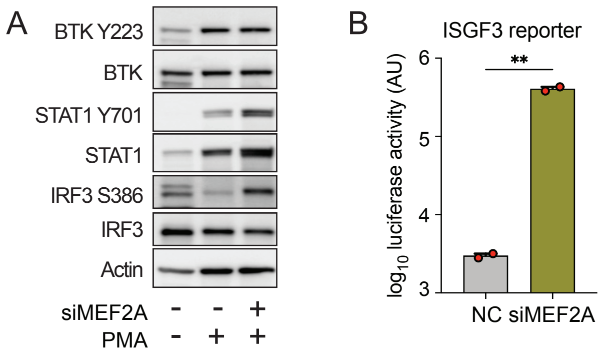


Supplementary Figure 5

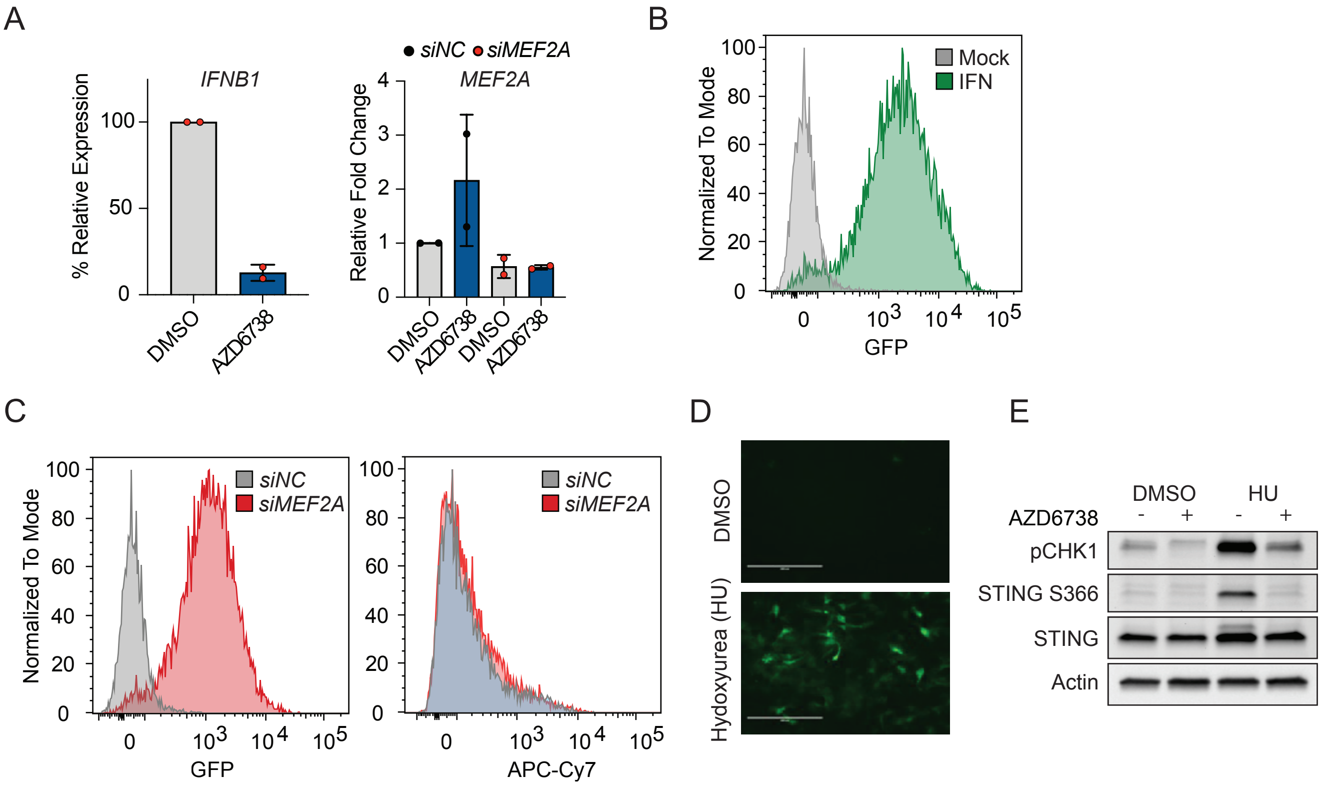


Supplementary Figure 6

| Gene_ID | FC.any.DE | pvalues |
|---|---|---|
| COLEC10 | -1.721494 | 9.88E-05 |
| ENPP4 | -1.2509886 | 1.27E-05 |
| MYZAP | -1.2306608 | 0.00032067 |
| CCDC173 | -1.2164092 | 0.00030939 |
| TNFRSF10D | -1.2120525 | 2.85E-05 |
| PTPN20 | -1.155903 | 0.00017895 |
| MIR600HG | -1.1001459 | 0.0003829 |
| LOC100147773 | -1.0829727 | 2.23E-05 |
| DANCR | -1.0812674 | 3.49E-05 |
| FILIP1 | -1.0805719 | 0.0002719 |
| C17orf104 | -1.0524818 | 0.00042748 |
| LOC148709 | -0.9705356 | 0.00041527 |
| LINC01600 | -0.9636841 | 0.00013035 |
| ACO1 | -0.9597223 | 2.75E-06 |
| IMMP1L | -0.9268096 | 0.00017374 |
| NDUFV2-AS1 | -0.9100478 | 0.00044377 |
| HTR1D | -0.8864776 | 0.00016405 |
| MTRNR2L2 | -0.8585358 | 0.00014181 |
| ZNF214 | -0.8021995 | 0.00030322 |
| FAM101B | -0.8004756 | 7.08E-06 |
| LOC100126784 | -0.7987506 | 0.00041209 |
| MTURN | -0.786232 | 0.0001048 |
| HSPA4L | -0.7652938 | 0.00026935 |
| KIAA0895 | -0.7612715 | 7.78E-05 |
| FAM172A | -0.7607289 | 0.00021323 |
| TMEM168 | -0.7478181 | 0.00043747 |
| GPATCH11 | -0.7440873 | 2.48E-05 |
| LAPTM5 | -0.7440816 | 0.00030389 |
| RRN3P2 | -0.7425545 | 0.00044862 |
| CERK | -0.7399386 | 4.35E-06 |
| LINC01094 | -0.737529 | 0.00024881 |
| ODC1 | -0.7225215 | 5.08E-05 |
| APPL2 | -0.7206027 | 1.28E-05 |
| GPAM | -0.7162097 | 0.00033311 |
| WRB | -0.7157394 | 8.74E-05 |
| SLC25A15 | -0.7106933 | 0.00028121 |
| MEGF9 | -0.7079503 | 4.56E-05 |
| HCG11 | -0.6919309 | 8.04E-05 |
| IL17RD | -0.6916286 | 0.00027462 |
| ANXA10 | -0.6894093 | 0.00042264 |
| GPCPD1 | -0.6807389 | 8.04E-05 |
| C8orf88 | -0.6750285 | 0.00017232 |
| NACC2 | -0.645078 | 0.00020281 |
| GATSL2 | -0.6355677 | 0.00017325 |

| | | |
|---|---|---|
| MTSS1 | -0.6275723 | 0.00044348 |
| SLC4A8 | -0.6264251 | 2.38E-05 |
| ACBD7 | -0.623978 | 0.00029793 |
| IPP | -0.6132339 | 0.00033407 |
| FAM81A | -0.6105034 | 7.31E-05 |
| NT5DC2 | -0.6030853 | 6.55E-05 |
| KATNAL2 | -0.6030397 | 0.0001939 |
| PDZD8 | -0.6017202 | 5.77E-05 |
| CMTM4 | -0.5992442 | 4.71E-05 |
| KRT80 | -0.5943656 | 8.85E-05 |
| HSPA12A | -0.5811236 | 8.89E-05 |
| ZNF788 | -0.5803003 | 0.0004044 |
| SLC12A6 | -0.5703899 | 0.0002186 |
| STRBP | -0.5702911 | 0.00012457 |
| AGAP1 | -0.5695165 | 5.34E-05 |
| MEF2A | -0.5652724 | 4.33E-05 |
| LOC646903 | -0.5633098 | 5.80E-05 |
| PLA2G12A | -0.5626649 | 0.00018495 |
| ADGRA3 | -0.5565492 | 0.00024908 |
| PLEKHM3 | -0.5518743 | 0.0003301 |
| GK | -0.5499953 | 0.00022969 |
| TNFAIP8L1 | -0.5396515 | 0.00017684 |
| CLN8 | -0.5377079 | 0.00023315 |
| TMEM107 | -0.5246044 | 4.19E-05 |
| SH3D19 | -0.5239952 | 0.00014807 |
| KHK | -0.5199502 | 0.00024265 |
| SLC7A2 | -0.5176353 | 9.63E-05 |
| TACC2 | -0.5145649 | 0.00011879 |
| LRP12 | -0.5125351 | 0.00028054 |
| TSPAN12 | -0.5111883 | 0.00011538 |
| PCOLCE2 | -0.5083587 | 0.00033572 |
| KATNAL1 | -0.5051265 | 0.00031264 |
| BRI3BP | -0.5047414 | 0.00039251 |
| MPHOSPH6 | -0.5033123 | 0.00031831 |
| PRKAR2A | -0.502514 | 5.72E-05 |
| SRSF8 | -0.502307 | 0.00013416 |
| CCDC15 | -0.4990238 | 0.00019981 |
| FAM168B | -0.4982571 | 3.28E-05 |
| APOLD1 | -0.4959591 | 0.00014509 |
| ZYG11B | -0.4949707 | 0.00021164 |
| KCTD7 | -0.4918453 | 5.98E-05 |
| E2F8 | -0.4911132 | 0.00016375 |
| GK5 | -0.4909303 | 6.90E-05 |
| TRIM45 | -0.4907377 | 0.00025111 |
| FAM216A | -0.4900525 | 0.00022393 |

| | | |
|---|---|---|
| SLC35G1 | -0.4888692 | 0.00030133 |
| PHKB | -0.4882101 | 0.00013832 |
| BUB1B | -0.4880397 | 5.16E-05 |
| SLC35D1 | -0.4803284 | 8.71E-05 |
| SETD7 | -0.4799788 | 0.00019643 |
| ATP1B1 | -0.4732573 | 0.00022329 |
| METTL7A | -0.4721484 | 6.20E-05 |
| FAM83D | -0.4686233 | 9.27E-05 |
| KCTD10 | -0.4679901 | 9.87E-05 |
| KIAA0232 | -0.4665354 | 9.78E-05 |
| TMX4 | -0.4643784 | 0.00037612 |
| UBE2D4 | -0.4627812 | 0.00037306 |
| TNFRSF9 | -0.4610672 | 0.00037995 |
| PXMP4 | -0.4609511 | 0.00021638 |
| NCAPD3 | -0.4590387 | 9.95E-05 |
| LOC730101 | -0.4566814 | 0.0001057 |
| CDKN3 | -0.4543003 | 0.00011032 |
| TFRC | -0.4539197 | 0.00019316 |
| PHF20 | -0.4521649 | 5.17E-05 |
| ZHX3 | -0.4520342 | 0.00016061 |
| H2AFV | -0.4517354 | 0.00013088 |
| PDSS1 | -0.4513158 | 0.00033901 |
| TIFA | -0.4499945 | 0.00015376 |
| USP13 | -0.4485907 | 0.00026965 |
| ASB1 | -0.4449271 | 0.0001295 |
| DDHD1 | -0.4445029 | 0.00034892 |
| BZW1 | -0.4437288 | 0.00017125 |
| NFYA | -0.4431595 | 8.79E-05 |
| MAN1A2 | -0.4410928 | 0.00037976 |
| PRR11 | -0.4404712 | 0.00017041 |
| VKORC1L1 | -0.4365751 | 0.00013281 |
| TRAK2 | -0.4324948 | 0.00026845 |
| RAB3B | -0.4309255 | 0.00011613 |
| HDGFRP3 | -0.4303089 | 0.00020657 |
| PHYH | -0.4295439 | 0.00023589 |
| ARHGEF39 | -0.4288398 | 0.0002613 |
| TMEM200B | -0.427306 | 0.00028754 |
| TIMM21 | -0.427068 | 7.01E-05 |
| FBXL4 | -0.4242447 | 0.00038918 |
| FOPNL | -0.4222555 | 0.00030669 |
| CDCA3 | -0.4221304 | 0.00012578 |
| PKNOX1 | -0.4194849 | 0.00015879 |
| CXXC5 | -0.4185443 | 0.00031707 |
| BTBD7 | -0.4178716 | 0.0002284 |
| SPAST | -0.4170873 | 0.0003392 |

| | | |
|---|---|---|
| HJURP | -0.4168782 | 8.80E-05 |
| CYP20A1 | -0.4162096 | 0.00033579 |
| SNX30 | -0.4106738 | 0.00026903 |
| C12orf49 | -0.4106472 | 0.00023485 |
| NBEA | -0.4097267 | 0.0003129 |
| UNC119B | -0.4083775 | 0.00011 |
| ABHD2 | -0.4080158 | 0.00011741 |
| MYO5A | -0.4059957 | 9.22E-05 |
| ZAK | -0.4032072 | 0.00028064 |
| PIK3IP1 | -0.402363 | 0.00012323 |
| CDCA8 | -0.4020812 | 0.0002791 |
| DNAJB5 | -0.401306 | 0.00042109 |
| C17orf58 | -0.4003422 | 0.00025292 |
| NAPB | -0.4002147 | 0.00031252 |
| HENMT1 | -0.4001638 | 0.00038756 |
| GSK3B | -0.4001253 | 0.00040546 |
| TRIM24 | -0.3946826 | 0.00043818 |
| C16orf72 | -0.3946556 | 0.00044489 |
| PRKCA | -0.3924311 | 0.00024609 |
| ITGAE | -0.3915823 | 0.00044031 |
| SIPA1L2 | -0.3913821 | 0.0003418 |
| EIF4EBP2 | -0.3908575 | 0.00028308 |
| PTPN14 | -0.3904806 | 0.00010075 |
| HAUS2 | -0.3903464 | 0.00037418 |
| PSRC1 | -0.3860963 | 0.00025381 |
| ZBTB8A | -0.3847996 | 0.00028622 |
| SORT1 | -0.382291 | 0.0002089 |
| CREBL2 | -0.3778023 | 0.00028201 |
| TBC1D14 | Spast | 0.00018792 |
| KNSTRN | -0.3748006 | 0.00013427 |
| PRC1 | -0.3745225 | 0.00011471 |
| PTPN11 | -0.3741294 | 0.00043726 |
| PABPC4 | -0.3679985 | 0.00035836 |
| TMEM241 | -0.3639387 | 0.00022931 |
| VAPB | -0.3626294 | 0.00043568 |
| CCNF | -0.3608864 | 0.00033003 |
| ZCCHC14 | -0.3594917 | 0.0001757 |
| CEP78 | -0.3587015 | 0.00025569 |
| CCNB2 | -0.3575143 | 0.00022337 |
| GCLM | -0.3564044 | 0.00034286 |
| SERTAD2 | -0.3561143 | 0.00034229 |
| AURKA | -0.3532296 | 0.00014344 |
| MSANTD3 | -0.3519649 | 0.00021512 |
| KLHL42 | -0.3506206 | 0.00044425 |
| METTL9 | -0.3499217 | 0.00038007 |

| | | |
|---|---|---|
| HIPK1 | -0.3497465 | 0.00041286 |
| AOX1 | -0.344831 | 0.00028469 |
| HACL1 | -0.3430892 | 0.00018423 |
| FAM64A | -0.3429971 | 0.00041699 |
| MAP7D3 | -0.341671 | 0.00042622 |
| RACGAP1 | -0.3413094 | 0.00032256 |
| TFDP2 | -0.3407587 | 0.00039029 |
| TPX2 | -0.3364613 | 0.00020747 |
| WDR12 | -0.3362257 | 0.00044202 |
| PRUNE | -0.3359956 | 0.00021981 |
| UBE4B | -0.335925 | 0.00042898 |
| RBMX | -0.3329687 | 0.0004337 |
| MRPL30 | -0.3213891 | 0.00029609 |
| BPNT1 | -0.320864 | 0.000357 |
| NXPE3 | -0.3116976 | 0.00032817 |
| TACC1 | -0.3108026 | 0.00044096 |
| PRICKLE2 | -0.3025744 | 0.00039892 |
| RCAN3 | -0.2961361 | 0.00044272 |
| TSPYL4 | -0.2958301 | 0.00030574 |
| DUSP3 | -0.2930827 | 0.00043693 |
| DOCK5 | -0.2927943 | 0.00040495 |
| LOC642852 | -0.291621 | 0.00039264 |
| PIK3R3 | 0.30320345 | 0.00040296 |
| IRS1 | 0.30600575 | 0.00026909 |
| PARD3 | 0.31416015 | 0.00031791 |
| RFX5 | 0.32290391 | 0.0004172 |
| BTN2A1 | 0.32300456 | 0.00043679 |
| CALCOCO2 | 0.3234099 | 0.00035796 |
| HSPA5 | 0.3364553 | 0.00035922 |
| CDK17 | 0.34481552 | 0.0002439 |
| KIAA0513 | 0.35043537 | 0.00024327 |
| SEMA3F | 0.36128487 | 0.00044574 |
| PTPRK | 0.36128795 | 0.00022612 |
| SPTLC2 | 0.36489426 | 0.00022905 |
| PFKFB4 | 0.37293881 | 0.00025069 |
| TBX15 | 0.37672198 | 0.00027692 |
| B4GALT5 | 0.37813639 | 0.0002132 |
| EPHA4 | 0.38097373 | 0.00018498 |
| MVP | 0.38229263 | 0.00032148 |
| DENND5A | 0.38560352 | 0.00012789 |
| PARP4 | 0.3993782 | 0.00041378 |
| TXNIP | 0.4019321 | 0.00015681 |
| ZMYND8 | 0.40545988 | 0.00012161 |
| STK10 | 0.40949724 | 0.00038975 |
| BAZ2A | 0.40989525 | 0.00028298 |

| | | |
|---|---|---|
| STAT6 | 0.41490246 | 0.00016835 |
| APBB2 | 0.41707207 | 0.00011255 |
| CHSY1 | 0.42000218 | 9.54E-05 |
| TNS1 | 0.4250337 | 0.00023511 |
| ATXN7 | 0.42564874 | 0.00015446 |
| DOK3 | 0.42688426 | 0.00032561 |
| TNC | 0.42726426 | 0.00034374 |
| ZFP36L1 | 0.42813594 | 0.00024425 |
| TLE4 | 0.42865749 | 0.00011623 |
| GPBP1 | 0.43817901 | 0.00030898 |
| ZFHX3 | 0.44353823 | 0.00041305 |
| AFF1 | 0.44447918 | 8.72E-05 |
| LGALS8 | 0.44579603 | 0.00026523 |
| ZNF710 | 0.45042252 | 0.0003301 |
| TMEM63A | 0.45083644 | 0.00034248 |
| ETV6 | 0.45104236 | 0.00011747 |
| PRKCD | 0.45248075 | 0.00013913 |
| DENND1A | 0.46797236 | 0.00038832 |
| TAF8 | 0.47022332 | 4.70E-05 |
| ENDOD1 | 0.47638275 | 0.00024987 |
| HIVEP1 | 0.47786235 | 0.00029078 |
| NUPR1 | 0.48014079 | 5.05E-05 |
| MICB | 0.48183478 | 9.78E-05 |
| PTN | 0.49113674 | 0.00012119 |
| PTGER2 | 0.49756348 | 9.98E-05 |
| CDC42EP3 | 0.4998019 | 0.00012735 |
| SPATA13 | 0.50001291 | 0.00021925 |
| SERPINE1 | 0.50127381 | 0.00024344 |
| LINC01000 | 0.50365961 | 5.88E-05 |
| GANC | 0.50513131 | 7.99E-05 |
| STX17 | 0.51039653 | 0.00028926 |
| HOXB4 | 0.51469789 | 0.00035886 |
| PSME1 | 0.51558537 | 9.77E-05 |
| SLC9A3R2 | 0.51681539 | 4.46E-05 |
| PDLIM1 | 0.51773436 | 7.26E-05 |
| DYNLT1 | 0.51813536 | 0.00011187 |
| RPS6KC1 | 0.52101692 | 0.00024283 |
| CPED1 | 0.52167522 | 0.00027593 |
| B2M | 0.52533275 | 0.00024123 |
| ITPRIPL2 | 0.52867223 | 2.53E-05 |
| AUTS2 | 0.528989 | 0.0002229 |
| EDARADD | 0.52958291 | 0.00011656 |
| RIPK1 | 0.53390573 | 7.13E-05 |
| VEGFC | 0.53465433 | 3.26E-05 |
| ERAP1 | 0.53560617 | 4.11E-05 |

| | | |
|---|---|---|
| GOLM1 | 0.53587182 | 4.86E-05 |
| BAK1 | 0.53930644 | 8.97E-05 |
| S1PR3 | 0.53962006 | 7.13E-05 |
| SGK1 | 0.54362235 | 4.79E-05 |
| DNAJA1 | 0.54381412 | 3.87E-05 |
| UHRF1BP1 | 0.54600687 | 0.00013555 |
| RAB24 | 0.54954319 | 0.00031084 |
| B3GALNT1 | 0.55129372 | 2.33E-05 |
| TLK2 | 0.55561932 | 1.76E-05 |
| VAMP5 | 0.55748738 | 0.00013402 |
| ITGA2 | 0.55815152 | 7.67E-05 |
| SCARB2 | 0.56265358 | 0.00013942 |
| DBF4B | 0.5639577 | 3.55E-05 |
| LIMA1 | 0.56400023 | 3.59E-05 |
| RRBP1 | 0.56517438 | 0.0002308 |
| USP42 | 0.56629679 | 4.88E-05 |
| OTUD4 | 0.56633663 | 3.11E-05 |
| FBXO32 | 0.57120093 | 3.80E-05 |
| ANKIB1 | 0.57169244 | 3.20E-05 |
| PLAUR | 0.57571283 | 0.00010175 |
| BTN3A2 | 0.57600111 | 2.50E-05 |
| SLC18B1 | 0.57609095 | 0.00040496 |
| RNF114 | 0.57692316 | 2.50E-05 |
| CD47 | 0.57731965 | 0.00018672 |
| MCL1 | 0.58079876 | 1.54E-05 |
| TBX20 | 0.58090904 | 0.0002855 |
| MAP3K14 | 0.58198062 | 3.13E-05 |
| PTGS2 | 0.58642757 | 0.00027158 |
| FNDC3A | 0.5877902 | 6.46E-05 |
| EPHB2 | 0.58851405 | 0.00033502 |
| PLCB4 | 0.58868388 | 0.00039708 |
| CDKL1 | 0.59070939 | 0.00031527 |
| RND3 | 0.59174597 | 2.93E-05 |
| C1RL-AS1 | 0.59435488 | 1.41E-05 |
| HDAC9 | 0.59576912 | 0.00011492 |
| PANX1 | 0.59767271 | 4.72E-05 |
| NR2F1 | 0.61519215 | 5.20E-05 |
| TAPBP | 0.62135822 | 9.91E-05 |
| GABRE | 0.62404308 | 0.00025694 |
| ERAP2 | 0.62459963 | 0.00018779 |
| YEATS2 | 0.62521788 | 4.10E-05 |
| OPTN | 0.62688872 | 1.45E-05 |
| FAM26E | 0.62749221 | 9.01E-05 |
| NOCT | 0.6320254 | 0.00016225 |
| NOD1 | 0.63274657 | 0.00031876 |

| | | |
|---|---|---|
| RHBDF2 | 0.63372105 | 5.05E-05 |
| PSORS1C1 | 0.63464414 | 0.00017106 |
| LINC01085 | 0.63659321 | 0.00013445 |
| TCF4 | 0.63692989 | 2.49E-05 |
| ZNF618 | 0.63825683 | 0.00011144 |
| RNF31 | 0.63904159 | 0.00010073 |
| SLC16A4 | 0.64222038 | 0.00019198 |
| TEC | 0.64992833 | 0.00020487 |
| TLR4 | 0.65144905 | 8.90E-05 |
| FZD5 | 0.65221861 | 0.00023998 |
| DNPEP | 0.65253633 | 6.98E-05 |
| PATL1 | 0.65303399 | 5.76E-06 |
| TGIF2 | 0.65674708 | 5.27E-05 |
| FMR1 | 0.65701062 | 0.00011592 |
| PARP8 | 0.65750294 | 4.07E-05 |
| BCL6 | 0.6612716 | 3.81E-05 |
| FHL3 | 0.66430777 | 6.38E-05 |
| MYCBP2 | 0.66506498 | 1.62E-05 |
| LGMN | 0.66769608 | 0.00028441 |
| NUB1 | 0.67225175 | 2.60E-05 |
| RICTOR | 0.67267597 | 0.00024784 |
| LIPA | 0.67726529 | 0.00013169 |
| BAZ1A | 0.67826365 | 2.85E-05 |
| CTNNBL1 | 0.68016528 | 2.30E-05 |
| HIRA | 0.6860269 | 1.53E-05 |
| KIAA1217 | 0.68871755 | 5.13E-06 |
| ZFP36L2 | 0.69356439 | 0.00031722 |
| CFLAR | 0.69618763 | 4.47E-05 |
| USP28 | 0.69676651 | 2.40E-05 |
| PROCR | 0.70036148 | 2.42E-05 |
| ABCD1 | 0.70491751 | 0.00024236 |
| CHMP5 | 0.70634931 | 3.46E-05 |
| KIAA0226 | 0.71257352 | 6.78E-06 |
| ANKFY1 | 0.71270649 | 9.42E-06 |
| TTC38 | 0.7199069 | 5.32E-05 |
| GSDMD | 0.72128749 | 0.00033211 |
| PDE4B | 0.72501789 | 0.00020352 |
| TNFAIP3 | 0.73100651 | 7.55E-05 |
| WHAMM | 0.73379387 | 5.76E-05 |
| ITPRIP | 0.73716141 | 6.74E-05 |
| CXorf38 | 0.73849997 | 8.08E-05 |
| SLFN5 | 0.73960363 | 1.88E-05 |
| USF1 | 0.74179796 | 1.03E-05 |
| CTGF | 0.74262904 | 1.00E-05 |
| MYH7B | 0.74546583 | 0.00037146 |

| | | |
|---|---|---|
| RAD9A | 0.74766746 | 6.12E-05 |
| NFE2L3 | 0.7498113 | 0.00011705 |
| MAP2 | 0.75022279 | 5.04E-06 |
| AMER1 | 0.75051163 | 0.0001236 |
| ARMCX1 | 0.77056239 | 4.78E-05 |
| PSME2 | 0.77166353 | 2.39E-05 |
| SLC8A1 | 0.77470903 | 1.42E-05 |
| SCO2 | 0.77787031 | 0.00011001 |
| GTPBP2 | 0.77828551 | 2.50E-05 |
| BTN3A3 | 0.78126258 | 3.11E-05 |
| RGMB | 0.78164295 | 3.14E-06 |
| CYTH1 | 0.78289207 | 1.80E-05 |
| MTMR11 | 0.78765237 | 0.00010739 |
| HLA-C | 0.78853712 | 7.17E-05 |
| CLMP | 0.79034182 | 1.16E-05 |
| NRSN2-AS1 | 0.79221755 | 3.87E-05 |
| C17orf67 | 0.79425108 | 0.00022345 |
| RAB20 | 0.79487366 | 0.00013561 |
| RBMS2 | 0.79812704 | 3.61E-06 |
| CFH | 0.79968467 | 0.00010005 |
| RNF149 | 0.80136109 | 5.20E-06 |
| CASP7 | 0.80162582 | 1.48E-05 |
| FST | 0.80281952 | 2.96E-05 |
| NUDCD1 | 0.80369013 | 0.00025681 |
| HLA-B | 0.80542413 | 2.78E-05 |
| SPATS2L | 0.80721536 | 2.33E-05 |
| N4BP1 | 0.8083204 | 7.44E-06 |
| PSMB8-AS1 | 0.81040783 | 0.00014752 |
| NPR3 | 0.81093995 | 5.20E-05 |
| SHOX2 | 0.81503637 | 0.00010432 |
| DCP1A | 0.8172463 | 7.45E-06 |
| GPR180 | 0.81887999 | 0.00018731 |
| BIRC3 | 0.82354753 | 0.00011542 |
| SLC2A5 | 0.82366647 | 6.23E-05 |
| CASP10 | 0.82849152 | 2.22E-05 |
| SOGA3 | 0.83154141 | 0.00010433 |
| CPEB3 | 0.83402106 | 0.0001071 |
| DDIT3 | 0.83863063 | 1.40E-05 |
| MFSD12 | 0.83883134 | 0.00018409 |
| ELF1 | 0.83984811 | 2.41E-05 |
| TMEM173 | 0.84070548 | 1.61E-05 |
| MOV10 | 0.84072579 | 1.98E-05 |
| TTC39B | 0.84106668 | 1.68E-05 |
| NFKBIZ | 0.84135006 | 0.00011293 |
| TMEM106A | 0.84238776 | 0.00016773 |

| | | |
|---|---|---|
| JAG1 | 0.84724492 | 1.97E-05 |
| CASP4 | 0.84792192 | 1.07E-05 |
| C1S | 0.84929116 | 2.15E-06 |
| KLF6 | 0.85322136 | 4.65E-06 |
| TRAFD1 | 0.85397685 | 4.18E-06 |
| KCNT2 | 0.85626566 | 0.00010367 |
| GRINA | 0.85711992 | 3.83E-05 |
| SLC1A3 | 0.86438987 | 0.00014405 |
| NAMPT | 0.86544798 | 0.00034233 |
| JADE2 | 0.87250208 | 2.34E-05 |
| IL18BP | 0.88171672 | 0.00025087 |
| BBC3 | 0.8835118 | 0.00019189 |
| PSMB10 | 0.88454968 | 7.87E-05 |
| SLFN12 | 0.89503149 | 9.99E-06 |
| SLC2A12 | 0.89513657 | 1.05E-05 |
| NAPA | 0.89534057 | 2.98E-05 |
| PI4K2B | 0.89712538 | 5.88E-05 |
| ADPRHL2 | 0.89742313 | 3.29E-05 |
| PLEKHF1 | 0.89947538 | 3.88E-05 |
| NCOA7 | 0.90006378 | 6.05E-06 |
| CDK18 | 0.90119859 | 0.00025532 |
| PDGFRL | 0.90178009 | 1.56E-06 |
| SEPT4-AS1 | 0.90326277 | 0.0004302 |
| BCL2L13 | 0.91251178 | 1.57E-06 |
| C1R | 0.91854269 | 1.29E-05 |
| BATF3 | 0.92383198 | 0.00036949 |
| TNFRSF14 | 0.93159347 | 0.00014822 |
| TSKU | 0.93319195 | 1.42E-05 |
| EDN1 | 0.93522571 | 9.35E-06 |
| ATXN7L1 | 0.93629376 | 0.0001625 |
| CMTR1 | 0.93689055 | 2.08E-06 |
| GTPBP1 | 0.93716536 | 4.68E-05 |
| IL15 | 0.94269319 | 0.00019915 |
| BTN3A1 | 0.94647791 | 3.68E-06 |
| NT5C3A | 0.94682893 | 0.00026457 |
| CCND1 | 0.94698151 | 9.31E-05 |
| ZBTB42 | 0.94699318 | 0.00021602 |
| LNPEP | 0.94837264 | 6.02E-06 |
| LOC101927027 | 0.9666926 | 3.54E-05 |
| C14orf159 | 0.97562199 | 2.52E-05 |
| VSIG10L | 0.97888034 | 0.00010536 |
| STC2 | 0.9819286 | 4.33E-06 |
| ZNF107 | 0.98523267 | 0.00011693 |
| DUSP5 | 0.98580391 | 3.27E-05 |
| TRIM26 | 0.98702439 | 8.33E-06 |

| | | |
|---|---|---|
| PHACTR4 | 0.99826141 | 1.26E-06 |
| FGF2 | 1.00332011 | 7.50E-05 |
| SHISA5 | 1.00896051 | 1.22E-05 |
| SDPR | 1.00940833 | 1.77E-06 |
| MT2A | 1.01344849 | 4.01E-05 |
| IKBKE | 1.01720441 | 4.63E-05 |
| CSF1 | 1.01880968 | 5.05E-06 |
| CPEB2 | 1.023111 | 0.00016101 |
| GPR158 | 1.02854243 | 0.00028314 |
| TTLL6 | 1.0309274 | 1.55E-05 |
| GBP2 | 1.03899243 | 4.72E-05 |
| ZFYVE26 | 1.04290193 | 3.84E-06 |
| SQRDL | 1.04635935 | 6.69E-05 |
| NEDD9 | 1.05120961 | 2.58E-06 |
| MASTL | 1.05742704 | 3.04E-06 |
| JAK2 | 1.05893311 | 1.30E-05 |
| EHD4 | 1.06990358 | 1.38E-05 |
| RBM43 | 1.08090248 | 7.56E-06 |
| CNP | 1.09337121 | 2.11E-06 |
| SPSB1 | 1.0979857 | 1.15E-05 |
| IFI30 | 1.11177949 | 6.17E-06 |
| IL12A | 1.11406128 | 0.00035545 |
| LOC153684 | 1.11533893 | 4.32E-05 |
| TREX1 | 1.11720623 | 8.94E-05 |
| HK2 | 1.13029464 | 2.04E-06 |
| FAM71F2 | 1.13188937 | 2.29E-05 |
| DENND3 | 1.1320461 | 4.59E-06 |
| PATL2 | 1.13559151 | 5.85E-05 |
| FZD4 | 1.13630722 | 1.40E-05 |
| FGF1 | 1.13636002 | 3.91E-05 |
| HOXD4 | 1.14005576 | 9.28E-05 |
| IRF2 | 1.1440835 | 1.34E-06 |
| PCGF5 | 1.14720926 | 0.00012762 |
| TRIM56 | 1.14843281 | 1.48E-05 |
| APOBEC3D | 1.15224466 | 1.98E-06 |
| HLA-E | 1.15891221 | 1.31E-06 |
| LYSMD2 | 1.15968475 | 0.00010758 |
| C21orf91 | 1.16323568 | 0.00039597 |
| MOB3C | 1.16649415 | 2.59E-05 |
| REC8 | 1.18327412 | 2.13E-06 |
| CTSS | 1.18453899 | 6.18E-05 |
| SP140L | 1.18932226 | 1.32E-06 |
| LOC554223 | 1.18968141 | 0.00016652 |
| PSMB8 | 1.19589696 | 1.16E-06 |
| ARHGAP27 | 1.21103047 | 7.31E-05 |

| | | |
|---|---|---|
| PHF11 | 1.21529509 | 2.05E-05 |
| SERPING1 | 1.21771672 | 8.80E-05 |
| RGS20 | 1.2191788 | 0.00043761 |
| STARD5 | 1.22356171 | 5.40E-05 |
| ASPHD2 | 1.23084081 | 7.02E-05 |
| SEMA3A | 1.24311236 | 1.59E-05 |
| CTRL | 1.24652342 | 0.00038766 |
| BLZF1 | 1.25779929 | 9.51E-07 |
| PMAIP1 | 1.26113491 | 7.91E-06 |
| PLEKHN1 | 1.26502584 | 0.0001815 |
| GATA3 | 1.29046808 | 4.87E-05 |
| SIX1 | 1.29314585 | 0.00038156 |
| PDCD1LG2 | 1.29693606 | 3.64E-06 |
| ZC3HAV1 | 1.30016632 | 1.99E-07 |
| AGRN | 1.30504064 | 0.00013226 |
| CD68 | 1.31670505 | 3.70E-06 |
| RBCK1 | 1.33714449 | 2.67E-05 |
| TGM2 | 1.34391958 | 1.87E-05 |
| TRIM69 | 1.3627705 | 1.37E-05 |
| TMEM62 | 1.36710325 | 4.38E-06 |
| TMEM171 | 1.39213867 | 4.70E-05 |
| IFITM2 | 1.39254352 | 3.25E-06 |
| SLC6A9 | 1.39653228 | 8.89E-06 |
| TRIL | 1.42875731 | 0.0003375 |
| RASGRP3 | 1.44470569 | 2.32E-07 |
| VCAM1 | 1.44486284 | 2.14E-05 |
| PRKD2 | 1.44943325 | 1.04E-05 |
| IRF9 | 1.45268235 | 9.90E-07 |
| CARD16 | 1.45679126 | 5.02E-05 |
| CASP1 | 1.47375805 | 2.77E-06 |
| NLRC5 | 1.53579605 | 2.10E-06 |
| CCL7 | 1.53675173 | 0.00010208 |
| SOX9 | 1.54397622 | 6.09E-07 |
| RARRES3 | 1.55543026 | 1.48E-05 |
| FAM65B | 1.56193435 | 4.44E-07 |
| GCH1 | 1.57796478 | 1.52E-06 |
| RNF19B | 1.57976765 | 1.72E-07 |
| ADAR | 1.58085407 | 1.11E-07 |
| LGALS3BP | 1.58188702 | 1.04E-06 |
| NT5E | 1.58934467 | 1.10E-07 |
| SCAMP1-AS1 | 1.63936767 | 2.74E-05 |
| ANKRD45 | 1.64812155 | 0.00010831 |
| SLC25A28 | 1.65053936 | 2.77E-07 |
| ZCCHC2 | 1.66867404 | 1.76E-06 |
| IRF1 | 1.67526985 | 5.01E-07 |

| | | |
|---|---|---|
| RNF213 | 1.68078434 | 8.82E-08 |
| CXCL16 | 1.70290999 | 4.37E-05 |
| CCL2 | 1.70959659 | 7.52E-08 |
| CFB | 1.71066587 | 8.11E-07 |
| CLDN23 | 1.71458669 | 0.00044815 |
| PPM1K | 1.71633007 | 1.97E-07 |
| PTGIR | 1.73239319 | 1.57E-05 |
| KIAA1755 | 1.73278754 | 8.09E-05 |
| EIF2AK2 | 1.74915445 | 2.51E-06 |
| GBP3 | 1.75224015 | 9.08E-08 |
| APOBEC3F | 1.81788272 | 1.34E-06 |
| PML | 1.831151 | 3.18E-06 |
| HLA-F | 1.83522802 | 1.40E-06 |
| KIAA2012 | 1.83548245 | 0.00014861 |
| PSMB9 | 1.83632115 | 2.06E-06 |
| APOL2 | 1.84656338 | 1.29E-07 |
| NLRP3 | 1.86592683 | 4.26E-05 |
| LY6E | 1.86899255 | 1.45E-05 |
| TRIM5 | 1.86903263 | 6.13E-07 |
| PLEKHG4B | 1.87101776 | 0.00012118 |
| TRIM38 | 1.88711776 | 9.21E-08 |
| OGFR | 1.89785037 | 1.72E-06 |
| PHACTR1 | 1.90220132 | 6.03E-05 |
| STAT2 | 1.91879175 | 5.57E-08 |
| FAM46A | 1.9195582 | 1.19E-06 |
| UBA7 | 1.91959026 | 9.17E-07 |
| MSX1 | 1.93098492 | 8.01E-05 |
| WARS | 1.94457589 | 1.25E-07 |
| DDX60 | 1.99451494 | 6.09E-06 |
| APOBEC3G | 2.00102567 | 5.88E-07 |
| LRRN3 | 2.00824666 | 2.55E-05 |
| SAMHD1 | 2.01062222 | 6.12E-08 |
| TAP2 | 2.01759903 | 5.76E-08 |
| LOC101929723 | 2.03678489 | 3.70E-08 |
| IFITM3 | 2.04036705 | 1.60E-06 |
| KLF4 | 2.05310727 | 2.10E-06 |
| NMI | 2.07000359 | 1.61E-07 |
| TLR3 | 2.07970066 | 2.94E-06 |
| SP100 | 2.08490059 | 3.00E-08 |
| UNC93B1 | 2.09371589 | 4.48E-05 |
| PLEKHA4 | 2.10143247 | 4.06E-07 |
| ATP10A | 2.10228111 | 1.47E-06 |
| SECTM1 | 2.14941918 | 7.92E-07 |
| GJD3 | 2.15027378 | 9.25E-05 |
| SEMA4D | 2.15909042 | 3.96E-05 |

| | | |
|---|---|---|
| ZNFX1 | 2.16715181 | 6.38E-08 |
| LOC100419583 | 2.1959315 | 6.19E-08 |
| LRRTM2 | 2.20603281 | 0.00017556 |
| IFI16 | 2.23188193 | 1.85E-07 |
| LAG3 | 2.24289298 | 2.95E-05 |
| APOL1 | 2.24411045 | 3.25E-07 |
| ATF3 | 2.27947716 | 2.29E-07 |
| C5orf56 | 2.30414812 | 1.08E-05 |
| HES4 | 2.30578776 | 0.00027431 |
| IL22RA1 | 2.34148046 | 1.71E-05 |
| TRIM25 | 2.35933062 | 8.54E-08 |
| LAP3 | 2.40037509 | 2.62E-08 |
| MYD88 | 2.40619499 | 1.51E-07 |
| APOL3 | 2.43427141 | 4.89E-08 |
| ZMYND15 | 2.43435346 | 4.81E-05 |
| TDRD7 | 2.45590898 | 2.62E-08 |
| IL12RB1 | 2.45943604 | 0.00026118 |
| PNPT1 | 2.47827156 | 1.43E-07 |
| PLSCR1 | 2.47906077 | 5.85E-08 |
| STAT1 | 2.48767248 | 1.56E-08 |
| C3AR1 | 2.49023035 | 9.84E-05 |
| TAP1 | 2.4956395 | 8.35E-08 |
| DTX3L | 2.50136646 | 1.16E-08 |
| MLKL | 2.56707023 | 7.39E-08 |
| GBP1 | 2.59612782 | 7.44E-09 |
| IL6 | 2.59987924 | 2.58E-06 |
| SOCS1 | 2.62976054 | 3.57E-05 |
| DHX58 | 2.67263034 | 2.91E-08 |
| C19orf66 | 2.67692714 | 2.24E-07 |
| IFIT5 | 2.68163239 | 1.11E-08 |
| UBE2L6 | 2.68806759 | 1.09E-07 |
| APOL6 | 2.68947661 | 1.17E-08 |
| RUFY4 | 2.69456688 | 9.61E-05 |
| TRIM21 | 2.73205579 | 6.77E-08 |
| IFI35 | 2.73337187 | 2.01E-07 |
| TRIM14 | 2.73950732 | 5.90E-08 |
| PARP9 | 2.75286428 | 2.12E-08 |
| PARP10 | 2.77330861 | 2.93E-06 |
| GMPR | 2.78619169 | 5.55E-08 |
| PARP12 | 2.80286796 | 4.30E-09 |
| ATP8A1 | 2.84371987 | 1.45E-05 |
| C18orf61 | 2.84371987 | 1.45E-05 |
| TYMP | 2.86355034 | 3.29E-05 |
| TRANK1 | 2.94561121 | 4.31E-08 |
| CD274 | 2.97098525 | 4.63E-06 |

| | | |
|---|---|---|
| SP110 | 2.97506701 | 1.07E-08 |
| IL15RA | 2.97628542 | 0.00015377 |
| SAMD9 | 2.98132686 | 5.42E-07 |
| IFNL1 | 3.02055811 | 5.06E-07 |
| TRAPPC3L | 3.02055811 | 3.09E-07 |
| TRIM22 | 3.04059909 | 9.41E-09 |
| USP30-AS1 | 3.06796855 | 0.00030422 |
| TMEM229B | 3.07681517 | 0.00042375 |
| PARP14 | 3.14709337 | 4.97E-08 |
| IRF7 | 3.16869153 | 3.45E-06 |
| LOC100133669 | 3.16925332 | 9.44E-05 |
| ADAP1 | 3.1878115 | 9.82E-05 |
| ODF3B | 3.19566817 | 1.25E-06 |
| THEMIS2 | 3.20718126 | 1.57E-08 |
| IFI44 | 3.24310611 | 1.15E-08 |
| HERC5 | 3.28175035 | 1.99E-08 |
| DDX60L | 3.2926545 | 9.92E-08 |
| SAMD9L | 3.31444671 | 7.30E-08 |
| HERC6 | 3.34857456 | 1.09E-08 |
| EPSTI1 | 3.38321357 | 2.38E-08 |
| IL4I1 | 3.38833122 | 2.38E-06 |
| CH25H | 3.3983789 | 3.50E-05 |
| TMEM140 | 3.42231093 | 5.47E-07 |
| SLC15A3 | 3.4226066 | 3.33E-07 |
| DSP | 3.4634866 | 2.41E-06 |
| CXCL9 | 3.54887894 | 7.68E-05 |
| XAF1 | 3.55279788 | 2.42E-08 |
| HELZ2 | 3.55290972 | 3.50E-07 |
| OAS3 | 3.56008906 | 1.68E-08 |
| PDZD2 | 3.5739916 | 8.66E-09 |
| BISPR | 3.6185498 | 7.94E-07 |
| ALK | 3.65439444 | 4.42E-07 |
| HRASLS2 | 3.66943108 | 7.86E-06 |
| LAMP3 | 3.74642083 | 0.00031749 |
| ANGPTL1 | 3.80193687 | 9.64E-07 |
| KLHDC7B | 3.81166591 | 2.05E-06 |
| ETV7 | 3.84133853 | 6.42E-08 |
| IFI6 | 3.85268635 | 3.03E-06 |
| CIITA | 3.87315678 | 0.0004107 |
| HSH2D | 3.87354722 | 4.49E-06 |
| DLL1 | 3.99293668 | 0.00033369 |
| IL7 | 4.15178706 | 0.00043086 |
| BCL2L14 | 4.199374 | 1.18E-05 |
| TRIM31 | 4.27472501 | 1.30E-06 |
| EXOC3L1 | 4.27506689 | 0.00012771 |

| | | |
|---|---|---|
| DDX58 | 4.32160961 | 4.28E-09 |
| CD38 | 4.37732068 | 0.00036779 |
| IFIH1 | 4.38841907 | 6.93E-09 |
| ISG20 | 4.49389754 | 4.96E-07 |
| CEACAM1 | 4.4994969 | 1.66E-05 |
| IFIT3 | 4.54833794 | 1.19E-09 |
| TNFSF10 | 4.55785456 | 1.32E-07 |
| FBXO6 | 4.66362128 | 4.30E-06 |
| CHRNA1 | 4.70049545 | 6.78E-05 |
| IFITM1 | 4.70898133 | 9.59E-09 |
| PIK3AP1 | 4.81624308 | 1.15E-07 |
| SSTR2 | 4.90881322 | 0.0001051 |
| ISG15 | 4.93129303 | 2.46E-08 |
| HCAR2 | 4.95813274 | 1.45E-08 |
| LMO2 | 4.97647332 | 0.00031924 |
| GRIP2 | 4.98314898 | 0.00012106 |
| MAB21L2 | 5.09194707 | 8.06E-05 |
| ZBP1 | 5.11223956 | 1.30E-06 |
| IFIT1 | 5.19791914 | 1.35E-09 |
| RTP4 | 5.21348142 | 2.06E-05 |
| LGALS9 | 5.26054917 | 3.85E-07 |
| TNFSF13B | 5.28660485 | 8.58E-05 |
| IFNB1 | 5.40533716 | 0.00042879 |
| OASL | 5.55329992 | 1.65E-07 |
| CACNA1I | 5.58982988 | 0.00016747 |
| GBP4 | 5.76687791 | 3.14E-07 |
| IFIT2 | 5.80070663 | 6.86E-10 |
| BATF2 | 5.80594181 | 7.23E-07 |
| IDO1 | 5.91170579 | 0.00045202 |
| C16orf47 | 5.94103866 | 3.49E-08 |
| GBP1P1 | 5.96286896 | 4.02E-05 |
| USP18 | 6.00330183 | 6.14E-09 |
| MX1 | 6.03296315 | 3.85E-09 |
| GBP5 | 6.04443237 | 3.06E-08 |
| CCL5 | 6.14614962 | 2.02E-07 |
| BST2 | 6.14909563 | 2.10E-06 |
| IFI27 | 6.37019774 | 2.66E-07 |
| CXCL11 | 6.57901385 | 2.08E-05 |
| OAS2 | 6.6651445 | 9.37E-11 |
| IFI44L | 6.84529093 | 3.59E-08 |
| PAX5 | 7.28347393 | 1.86E-09 |
| OAS1 | 7.34344418 | 8.62E-10 |
| CXCL10 | 8.44807765 | 5.24E-05 |
| CMPK2 | 8.93662242 | 1.34E-06 |
| MX2 | 9.11564358 | 4.63E-09 |

| | | |
|---|---|---|
| RSAD2 | 10.0262142 | 1.91E-05 |

**Key resources table**

| REAGENT or RESOURCE | SOURCE | IDENTIFIER |
|---|---|---|
| **Antibodies** | | |
| Donkey Anti-Mouse IgG (HRP-Conjugated) | Jackson ImmunoResearch | RRID: AB_2340770 |
| Donkey Anti-Rabbit IgG (HRP-Conjugated) | Jackson ImmunoResearch | RRID: AB_10015282 |
| b-Actin-HRP (13E5) | Cell signaling | RRID: AB_1903890 |
| Mouse anti DNA-RNA hybrid [S9.6] | Kerafast | RRID:AB_2687463 |
| Rabbit anti GAPDH (D16H11) | Cell signaling technology | RRID:AB_10622025 |
| Rabbit anti cGAS (D1D3G) | Cell signaling technology | RRID:AB_2799712 |
| Rabbit anti MEF2A | Bethyl laboratories | RRID:AB_10954235 |
| Rabbit anti STAT1 | Cell signaling technology | RRID:AB_2197984 |
| Rabbit anti phosphorylated STAT1 (Y701) | Cell signaling technology | RRID:AB_561284 |
| Rabbit anti IRF1 (D5E4) | Cell signaling technology | RRID:AB_10949108 |
| Rabbit anti IRF3 (D6I4C) | Cell signaling technology | RRID:AB_2722521 |
| Rabbit anti IRF3 (phospho S386) [EPR2346] | Abcam | RRID:AB_1523836 |
| Rabbit anti DDX41 | Thermo Scientific | RRID:AB_2809226 |
| Mouse anti IFI16 | Santa Cruz | RRID:AB_627775 |
| Rabbit anti p-STING | Cell signaling technology | RRID:AB_2737062 |
| Rabbit anti STING | Cell signaling technology | RRID:AB_2799947 |
| Rabbit anti phospho histone H2A.X (Ser139) | Cell signaling technology | RRID:AB_2118010 |
| Rabbit anti ATF4 (D4B8) | Cell signaling technology | RRID:AB_2616025 |
| Rabbit anti XBP-1s (D2CF1F) | Cell signaling technology | RRID:AB_2687943 |
| Rabbit anti pCHK1 | Cell signaling technology | RRID:AB_331212 |
| Rabbit anti phospho RPA32 | Bethyl laboratories | RRID:AB_2180847 |
| Rabbit anti RPA32 | Cell Signaling Technologies | RRID:AB_2238543 |
| **Bacterial and virus strains** | | |
| VSV-GFP | Michael Gale | Fredericksen and Gale, 2006 |
| Coxsackie Virus B3 (Nancy) | Raul Andino | Laufman et al., 2019 |
| Sendai Virus (Cantell) | Charles River Lab | |
| Stellar Competent Cells *E. coli* HST08 strain | Takara | Cat# 636763 |
| **Chemicals, peptides, and recombinant proteins** | | |
| Human recombinant IFNb | PBL Assay Science | 111415-1 |
| Human recombinant IFNl3 | R&D Systems | 5259-IL-025 |
| Mirus Bio TransIT-X2 | Fisher Scientific | MIR6000 |
| Mirus Bio TransIT-TKO | Fisher Scientific | MIR2150 |
| DAPI | Thermo Fisher Scientific | D1306 |
| RNase T1 | Thermo Fisher Scientific | EN0541 |

| | | |
|---|---|---|
| RNase III | Thermo Fisher Scientific | AM2290 |
| RNase H | Thermo Fisher Scientific | AM2292 |
| B18R | Invitrogen | 34-8185-81 |
| Etoposide | Sigma-Aldrich | 341205 |
| Thapsigargin | Sigma-Aldrich | T9033 |
| Nocodazole | Sigma-Aldrich | M1404 |
| ATR inhibitor, ETP-46465 | Cayman Chemicals | 19809 |
| ATR inhibitor, AZD6738 | Cayman Chemicals | 21035 |
| ATM inhibitor, KU-55933 | SelleckChem | S1092 |
| DNA-Pk inhibitor, NU7441 | SelleckChem | S26638 |
| TBK1 inhibitor, GSK8612 | SelleckChem | S8872 |
| Hydroxyurea | Sigma-Aldrich | H8627-1G |
| 100x Halt protease and phosphatase inhibitor | Thermo Fisher Scientific | 78420 |
| Ibrutinib | Med Chem Express | PCI-32765 |
| **Critical commercial assays** | | |
| NucleoSpin RNA II | Macherey-Nagel | 740955.25 |
| QuantiTect RT kit | QIAGEN | 205314 |
| iSCRIPT cDNA Synthesis Kit | Bio-Rad | 1708891 |
| SsoAdvanced Universal Probes Supermix | Bio-Rad | 1725281 |
| TaqMan Universal Master Mix II, no UNG | Thermo Fisher Scientific | 4440048 |
| TruSeq Stranded mRNA Library Prep Kit | Illumina | 20020594 |
| Qubit RNA BR Assay Kit | Thermo Fisher Scientific | Q10210 |
| Pierce Gaussia Luciferase Glow Assay Kit | Thermo Fisher Scientific | 16160 |
| NucleoSpin Tissue DNA kit | Macherey-Nagel | 740952.5 |
| Pierce BCA Protein Assay | Thermo Fisher Scientific | PI23227 |
| CelLytic NyCLEAR Extraction Kit | Millipore Sigma | NXTRACT-1KT |
| 2',3'-Cyclic GAMP ELISA Kit | Arbor Assays | K067-H1 |
| Qubit RNA BR assay kit | Invitrogen | Q10210 |
| RNA 6000 Nano Kit | Agilent | 5067-1511 |
| **Deposited data** | | |
| RNA sequencing data | This paper | GEO: GSE209601 |
| **Experimental models: Cell lines** | | |
| AC16 | Millipore | Cat# SCC109, RRID:CVCL_4U18 |
| BJ/TERT | Saumendra Sarkar | RRID:CVCL_6573 |
| U937 | ATCC | Cat# CRL-1593.2, RRID:CVCL_0007 |
| THP-1 | ATCC | Cat# TIB-202, RRID:CVCL_0006 |
| Huh7 | ATCC | RRID: CVCL_0336 |
| *TMEM173* KO AC16 | This Paper | |
| *IRF3* KO AC16 | This Paper | |
| *CGAS* KO AC16 | This Paper | |
| *IFI16* KO AC16 | This Paper | |

| | | |
|---|---|---|
| *DDX41* KO AC16 | This Paper | |
| 5xISGF3-GLuc Huh7 reporter cells | This Paper | |
| pISRE-sfGFP AC16 | This Paper | |
| **Oligonucleotides** | | |
| DsiRNA MEF2A | Integrated DNA technologies | hs.Ri.MEF2A.13.2 |
| DsiRNA MEF2A | Integrated DNA technologies | hs.Ri.MEF2A.13.1 |
| siRNA MEF2A | Thermo Fisher Scientific | Assay ID 107760 |
| DsiRNA Negative control | Integrated DNA technologies | 51-01-14-04 |
| siRNA Negative control | Thermo Fisher Scientific | AM4613 |
| DsiRNA TMEM173 | Integrated DNA technologies | hs.Ri.TMEM173.13.1 |
| DsiRNA MAVS | Integrated DNA technologies | hs.Ri.MAVS.13.1 |
| DsiRNA MEF2C | Integrated DNA technologies | hs.Ri.MEF2C.13.2 |
| DsiRNA MEF2D-1 | Integrated DNA technologies | hs.Ri.MEF2D.13.1 |
| DsiRNAMEF2D-2 | Integrated DNA technologies | hs.Ri.MEF2D.13.2 |
| siRNA IRF1 | Thermo Fisher Scientific | 115266 |
| siRNA STAT1 | Fisher Scientific | M-003543-01-0005 |
| MEF2A | Life Technologies | Hs01050406_g1 |
| MEF2B | Integrated DNA technologies | Hs.PT.58.39043087 |
| MEF2C | Integrated DNA technologies | Hs.PT.58.14426705 |
| MEF2D | Integrated DNA technologies | Hs.PT.58.20959128 |
| IFNB1 | Integrated DNA technologies | Hs.PT.58.39481063.g |
| CXCL10 | Thermo Fisher Scientific | Hs00171042_m1 |
| ISG15 | Integrated DNA technologies | Hs.PT.58.39185901.g |
| HPRT1 | Integrated DNA technologies | Hs.PT.58v.45621572 |
| CVB3 VP1<br>F: 5′-ACGAATCCCAGTGTGTTTTGG-3′<br>R: 5′-TGCTCAAAAACGGTATGGACAT-3′ | Integrated DNA technologies | |
| CHMP2A<br>F: 5'-CGCGAGCGACAGAAACTAGAG-3'<br>R: 5'-CCCGCATCAATACAAACTTGC-3' | Integrated DNA technologies | |
| 5xISGF3_BS-hGLuc_PEST gBLOCK<br>5'-<br>ATCTCGATCGAAGAAATGAAACTTGATCATCGCCGAAGAAA<br>TGAAACTGCGAATCTGACGAAGAAATGAAACTCGCTTCGTA<br>ACGAAGAAATGAAACTCGTAGACTACCGAAGAAATGAAACT<br>CCCGGGTAGGGCCCAATTCGAGTCGAGGTAGGCGTGTAC<br>GGTGGGAGGTCTATATAAGCAGAGCTGGTTTAGTGAACCG<br>TCAGATCGCCTGGAGAGATCTTTGTCGATCCTACCATCCAC<br>TCGACACACCCGCCAGCGACCACTGCCAAGCTTCCGAGCT<br>CTCGCTCTAGAgccgccaccATGGGAGTCAAAGTTCTGTTTGC<br>CCTGATCTGCATCGCTGTGGCCGAGGCCAAGCCCACCGA<br>GAACAACGAAGACTTCAACATCGTGGCCGTGGCCAGCAAC<br>TTCGCGACCACGGATCTCGATGCTGACCGCGGGAAGTTGC | Integrated DNA technologies | |

| | | |
|---|---|---|
| CCGGCAAGAAGCTGCCGCTGGAGGTGCTCAAAGAGATGG AAGCCAATGCCCGGAAAGCTGGCTGCACCAGGGGCTGTC TGATCTGCCTGTCCCACATCAAGTGCACGCCCAAGATGAA GAAGTTCATCCCAGGACGCTGCCACACCTACGAAGGCGAC AAAGAGTCCGCACAGGGCGGCATAGGCGAGGCGATCGTC GACATTCCTGAGATTCCTGGGTTCAAGGACTTGGAGCCCA TGGAGCAGTTCATCGCACAGGTCGATCTGTGTGTGGACTG CACAACTGGCTGCCTCAAAGGGCTTGCCAACGTGCAGTGT TCTGACCTGCTCAAGAAGTGGCTGCCGCAACGCTGTGCGA CCTTTGCCAGCAAGATCCAGGGCCAGGTGGACAAGATCAA GGGGGCCGGTGGTGACAAGCTCCCTAGAAGTCATGGATTC CCACCTGCAGTGGCCGCGCAGGACGATGGTACCCTACCG ATGTCTTGCGCTCAAGAGAGCGGAATGGACCGACATCCAG CGGCGTGTGCCTCAGCAAGAATTAACGTTTAG-3' | | |
| 5xISGF3_BS GA fwd 5'-ATTTTATTATCTAACTGCTGATCGAGTGTAGCCAGATCTCC CGGGATCTCGATCGAAGAAATGAAACTTGATCA-3' | Integrated DNA technologies | |
| hLuc_PEST GA rev 5'-CTGTATTGCTACTTGTGATTGCTCCATGTTTTTCTAGGTCTC GAGCTAAACGTTAATTCTTGCTGAGGCACAC-3' | Integrated DNA technologies | |
| **Recombinant DNA** | | |
| pISRE-sfGFP | Nicholas Heaton | Froggatt et al., 2021 |
| pRRL-H1-PURO | Daniel Stetson | Gray et al., 2015 |
| pRRL-STING-PURO | Daniel Stetson | Gray et al., 2015 |
| pRRL-cGAS-PURO | Daniel Stetson | Gray et al., 2015 |
| pRRL-IRF3-PURO | Ram Savan | Schwerk et al., 2019 |
| pRRL-IFI16-PURO | Daniel Stetson | Gray et al., 2015 |
| pRRL-DDX41-PURO | This paper | |
| pTRIPZ-5xISGF3_BS-hGLuc_PEST | This paper | |
| **Software and algorithms** | | |
| FIJI | Schindelin et al., 2012 | https://imagej.net/software/fiji/ |
| CellProfiler | Sterling et al., 2021 | https://cellprofiler.org/ |
| GraphPad Prism 9 | GraphPad Software | RRID: SCR_002798 |
| R Studio 1.1.442 | Rstudio | RRID: SCR_000432 |
| FastQC (version 0.11.3) | http://www.bioinformatics.babraham.ac.uk/projects/fastqc/ | RRID: SCR_014583 |
| Bowtie2 (version 2.3.4) | Langmead and Salzberg, 2012 | RRID: SCR_016368 |
| STAR (version 2.5.3a) | Dobin et al., 2013 | RRID: SCR_004463 |
| HTSeq (version 0.6.1) | http://htseq.readthedocs.io/en/release_0.9.1/ | RRID: SCR_005514 |
| edgeR (version 3.20.9) | Robinson et al., 2010 | RRID: SCR_012802 |
| limma (version 3.34.8) | Ritchie et al., 2015 | |
| EnrichR | Chen et al., 2013 | RRID: SCR_001575 |
